# *DCP2* plays multiple roles during *Drosophila* development – possible case of moonlighting?

**DOI:** 10.1101/830729

**Authors:** Rohit Kunar, Jagat K Roy

## Abstract

mRNA decapping proteins (DCPs) are components of the P-bodies in the cell which are hubs of mRNAs targeted for decay and they provide the cell with a reversible pool of mRNAs in response to cellular demands. The *Drosophila* genome codes for two decapping proteins, DCP1 and DCP2 out of which DCP2 is the cognate decapping enzyme. The present endeavour explores the endogenous promoter firing, transcript and protein expression of *DCP2* in *Drosophila* wherein, besides a ubiquitous expression across development, we identify active expression paradigm during dorsal closure and a plausible moonlighting expression in the Corazonin neurons of the larval brain. We also demonstrate that the ablation of *DCP2* leads to embryonic lethality and defects in vital morphogenetic processes whereas a knockdown of *DCP2* in the Corazonin neurons reduces the sensitivity to ethanol in adults, thereby ascribing novel regulatory roles to DCP2. Our findings unravel novel putative roles for DCP2 and identify it as a candidate for studies on the regulated interplay of essential molecules during early development in *Drosophila*, nay the living world.

## Introduction

Organismal development mimics an orchestra with precisely timed and fine-tuned role(s) for each of the players. Balanced expression of genes requires timed activity of gene promoters at the proper site along with orderly degradation of transcripts and/or proteins (Yao and Ndoja, 2012; Ding, 2015). Decay of transcripts is one of the strategies to regulate gene expression (Ghosh and Jacobson, 2010) and the mRNA decapping proteins (DCPs) assume prime importance therein. These proteins initiate degradation of the mRNAs in cytoplasmic foci known as P-bodies, by removal of the 7-methylguanosine cap at the 5’ end of the mRNAs (Coller and Parker, 2004). The *Drosophila* genome codes for two mRNA decapping proteins, *viz.*, DCP1 and DCP2, out of which DCP2 is the cognate decapping protein. While DCP1 functions to activate DCP2 (Ren et al, 2012) and P-bodies/DCP1 have been implicated in miRNA mediated gene silencing (Rehwinkel et al, 2005), localization of the oskar mRNA in the *Drosophila* oocyte (Lin et al, 2006) and in oogenesis (Lin et al, 2008), DCP2 has been implicated in chronic nicotine induced locomotor hyperactivity in *Drosophila* (Ren et al, 2012). However, characterization of the role of decapping proteins in development has been limited to *Arabidopsis* (Xu et al, 2006) and *Caenorhabditis elegans* (Lall et al, 2005) only. Despite being the cognate decapping protein in *Drosophila*, the spatio-temporal dynamics of *DCP2* activity remains unexplored. The gene in *Drosophila* is ~8kb in length, has two curated promoters, *viz.*, a proximal promoter, DCP2_1 and a second, downstream promoter, DCP2_2 (Eukaryotic Promoter Database, EPD; Dreos et al, 2014) and codes for four transcripts (FlyBase; Drysdale, 2008). Herein, we have tried to explore the temporal activity of the *DCP2* promoter using the conventional UAS-GAL4 system (Brand and Perrimon, 1993) wherein, we used a GAL4 driven by the *DCP2* promoter (*DCP2*^*GAL4*^; Lukacsovich et al, 2001; Ren et al, 2012) and combined it with a modified UAS line (G-TRACE; Evans et al, 2009) to delineate the real-time promoter activity of *DCP2* during embryonic dorsal closure and in the larval tissues. In parallel, we endeavored to delineate the expression of the transcript isoforms or splice variants generated, and the expression paradigm of the translated protein. Although, *DCP2* is highly active and ubiquitous, high expression of the DCP2 protein was observed in the Corazonin (Crz) neurons of the larval brain and renders the individuals less sensitive to ethanol when knocked down in the corazonin neurons and it is expressed along the A-P and D-V axes in the larval wing pouch. *Loss-of-function* mutants of *DCP2* are embryonic lethal and showed defects in epithelial morphogenesis and organization of the embryonic nervous systems along with elevation and spatial disruption of the JNK cascade. Collectively, our observations present us with a stage for further exploration of hitherto undescribed facets of *DCP2* activity and identify DCP2 as a potential candidate for explication of molecular interplay during *Drosophila* development.

## Materials and Methods

### Fly strains, genetics and lethality assay

All flies were raised on standard agar-cornmeal medium at 24±1°C. *Oregon* R^+^ was used as the wild type control. For targeted gene expression (Brand and Perrimon, 1993), *DCP2*^*BG01766*^/*TM3,Ser*^1^ (*DCP2*^*GAL4*^; Ren et al, 2012), *CCAP-GAL4, TH-GAL4, Ap-GAL4, Ddc-GAL4, UAS-GFP, UAS-mCD8::GFP* and *UAS-DCP2*^*RNAi*^ were obtained from the Bloomington *Drosophila* Stock Centre, while G-TRACE (Evans et al, 2009) was a kind gift from Prof. Utpal Banerjee, MBI, UCLA. *DCP2^e00034^*/*TM3, Ser*^*1*^ was obtained from the Harvard *Drosophila* Stock Centre. The JNK signaling bio-sensor, *TRE-RFP* (Chatterjee and Bohmann, 2012; referred to as TRE-JNK in the text) was obtained as kind gift from Dr. B. J. Rao, TIFR, Mumbai, India while *sNPF-GAL4*, *Dilp2-GAL4* and *Crz-GAL4/CyO* were procured from Prof. Gaiti Hasan, NCBS, Bangalore, India.

*DCP2*^*BG01766*^/*TM3, Ser*^*1*^ and *DCP2*^*e00034*^/*TM3, Ser*^*1*^ were further introgressed with *TM3, ActGFP, Ser*^*1*^/TM6B stock in order to generate *DCP2*^*BG01766*^/*TM3, ActGFP, Ser*^*1*^ or *DCP2*^*e00034*^/*TM3, ActGFP, Ser*^*1*^ stocks. *TRE-JNK* (Chatterjee and Bohmann, 2012) was introgressed with *Sp/CyO; DCP2*^*BG01766*^/*TM3, ActGFP, Ser*^*1*^ or and *Sp/CyO; DCP2*^*e00034*^/ *TM3, ActGFP, Ser*^*1*^ to obtain *TRE-JNK*; *DCP2*^BG01766^/*TM3, ActGFP, Ser*^*1*^ and *TRE-JNK*; *DCP2*^*e00034*^/*TM3, ActGFP, Ser*^*1*^ stocks, respectively.

For behaviour assays, *Crz-Gal4/CyO* flies were crossed to *w^1118^* or *UAS-DCP2^RNAi^* flies to generate *Crz-Gal4/+* (Control) or *Crz-Gal4*/+; *UAS-DCP2*^*RNAi*^/+ (Experimental) genotypes.

Embryonic lethality was calculated as described in Bhuin and Roy, 2009. 100 embryos were transferred to agar plates and incubated for 24 to 48 h at 23°C and the total number of dead embryos was counted against total number of fertilized eggs. These fertilized eggs include the dead as well as the hatched embryos. The percentage of lethality was calculated as –

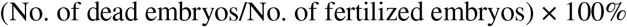

The percentage (%) lethality for each cross was calculated in triplicates and the mean lethality so obtained was tabulated and graphically represented using MS-Excel-2010. The final percentages have been calculated by multiplying the lethality obtained in every cross scheme with the inverse of the fraction of the progeny determined by standard Mendelian ratios.

### Detection of DCP2 transcript expression and analysis of splice variants

Detection of transcript expression from *DCP2* was performed by reverse-transcriptase polymerase chain reaction (RT-PCR) using a combination of primers designed such that the amplicon size would discriminate between the individual isoforms which included a single reverse primer, DCP2_DBAE_R, which could bind to all of the transcripts, and two forward primers, *viz.*, DCP2_BAE_F which would bind to isoforms DCP2-RA, RB and RE, and DCP2_D_F, which would bind to DCP2-RD. Being similar in architecture, DCP2-RA and RE would yield similar sized amplicons with the above primer pair, but DCP2-RD would yield a smaller amplicon. However, the 3’UTR is longer and unique for DCP2-RE and we exploited the architectural bias to discriminate between the two isoforms by designing an additional primer pair which would amplify the 3’UTR sequence of DCP2-RE uniquely. The table below (**Table 1**) shows the primer sequences, the combinations and the calculated amplicon sizes for each of the isoform with each of the primer pairs. The unique amplicons are italicized. RT-PCR was performed according to Lakhotia et al, 2012 in the samples discussed in the results section.

**Table 1:**
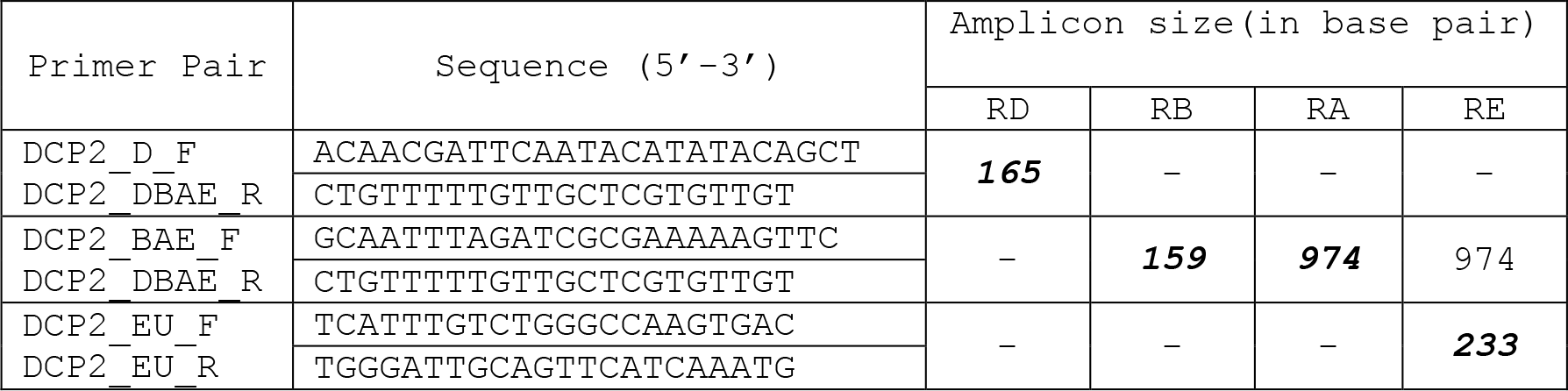
List of primer sequences, combinations generated and calculated amplicon sizes for detection of expression of *DCP2* transcript isoforms

### Embryo collection and fixation

All flies were made to lay eggs on standard agar plates supplemented with sugar and propanoic acid and eggs were collected according to Narasimha and Brown, 2006, with slight modifications. For whole mount preparations and immunostaining of embryos, different alleles and transgenes were balanced with GFP tagged balancers and only non-GFP or driven embryos were selected for experimental purpose. Embryo staging was done according to Hartenstein’s Atlas of *Drosophila* Development, 1993.

### Immunocytochemistry

*Drosophila* embryos were fixed and imaged as described by Narasimha and Brown, 2006. The dechorionated and devitellized embryos were fixed in 4% para-formaldehyde solution and stored in absolute methanol. Immunostaining of the embryos was done as described in Nandy and Roy, 2019. Late third instar larval tissues were dissected out in 1× PBS, fixed in 4% paraformaldehyde for 20 min at RT and immunostained as described previously in Banerjee and Roy, 2017. The primary antibodies used were – mouse anti-DCP2 (1:50; PCRP-DCP2-1D6, DSHB), mouse anti-Fasciclin II (1:100; 1D4, DSHB), mouse anti-Fasciclin III (1:100; 7G10, DSHB) and rabbit anti-phospho-JNK/SAPK (1:100; 81E11 Cell Signaling Technology). Appropriate secondary antibodies conjugated either with Cy3 (1:200, Sigma-Aldrich, India) or Alexa Fluor 488 (1:200; Molecular Probes, USA) or Alexa Fluor 546 (1:200; Molecular Probes, USA) were used to detect the given primary antibody, while chromatin was visualized with DAPI (4’, 6-diamidino-2-phenylindole dihydrochloride, 1μg/ml Sigma-Aldrich). For imaging of live embryos for real-time promoter analysis using G-TRACE or JNK signaling, embryos of the desired genotype were rinsed in 1X PBS, dechorionated in bleach and mounted in halocarbon oil and observed directly.

### Cuticular preparations from embryos

Cuticle preparations were made from embryos as described by Wieschaus and Nusslein-Volhard, 1986 along with some modifications as described in Sasikumar and Roy, 2008. Briefly, the eggs were dechorionated in bleach and washed in an aqueous solution containing 0.7% NaCl and 0.02% Triton-X 100. The eggs were washed thrice in 0.1% Triton-X, devitellinised in a mixture of methanol and n-heptane (1:1 v/v) mixture. They were fixed in 1 part glycerol - 4 parts acetic acid for 1 h, mounted in Hoyer’s mountant and cleared at 60°C overnight.

### Microscopy and Documentation

The immunostained slides were observed under Zeiss LSM 510 Meta Laser Scanning Confocal microscope, analysed with ZEN12 and LSM softwares and assembled using Adobe Photoshop 7.0. The cuticles were observed in dark field or phase-contrast optics, namely 10× Plan Fluor Ph1 DLL (0.3 NA), 20× Plan Fluor Ph1 DLL (0.5 NA) and 40× Plan Fluor Ph2 DLL (0.75 NA) objectives (Nikon, Japan) and the images were captured with a Nikon Digital camera DXM1200. Fluorescence imaging of embryos for analysis of *DCP2* promoter using G-TRACE or JNK signaling was done in Nikon 90i Fluorescence microscope under 10X Plan Fluor Ph1 DLL (0.3 NA), 20X Plan Fluor Ph1 DLL (0.5 NA) objectives.

### Behaviour Assays

Groups of 20 males and females (1-3 days old) of the desired genotypes, *viz., Crz-Gal4*/+ (Control) and *Crz-Gal4*/+; *UAS-DCP2*^*RNAi*^/+, maintained on food vials in a 12L: 12D conditions at 23°C for 1 day were used for the following behaviour assays.

### Ethanol Sedation Assay

Ethanol sedation assays were performed as described previously (McClure et al, 2013) with minor alterations. Briefly, flies were transferred to empty vials, sealed with cotton plugs and allowed to acclimatize for 10-20 min. The cotton plugs were replaced with fresh plugs containing 1 ml of 100% ethanol. They were maintained in such “booze chambers” for 15-20 mins. During the treatment, files were mechanically stimulated by tapping and/or mechanically swirling the vials at intervals of 5 mins. Flies able to climb the walls and/or move their appendages on the floor of the vial were considered “non-sedated” while those unable to execute such activity were considered “sedated”. The number of sedated flies was counted at 5 min intervals. The time to 50% sedation (ST50) was determined by manual extrapolation.

### Recovery from Ethanol Sedation

Recovery from ethanol induced sedation was assayed as described by Sha et al, 2014. Following exposure to ethanol (described above), the cotton plugs were replaced with fresh plugs and the vials were inverted to place them upside down. The number of “non-sedated” flies (considered as “recovered”) was counted every 10 min.

## Results and Discussion

### *DCP2* mRNAs are expressed ubiquitously throughout *Drosophila* development

In order to determine the presence or absence of DCP2 at a particular stage alongwith identification of the exact isoform(s)/splice variant(s) expressed therein, RT-PCR was performed using primers designed for the same. DCP2 expression was detectable at all stages of development, *viz.* embryo (0-24h), larval stages (1^st^, 2^nd^ and 3^rd^ instars), pupal and adult stages. Among the four annotated variants, DCP2_RE (FBtr0304975) and DCP2_RA (FBtr0075538) was observed in all stages of fly development, whereas DCP2_RD (FBtr0100528) was observed only in the larval gonads, *viz.* testes and ovaries, and in the adult flies (**Figure 1**). The other variant, DCP2_RB (FBtr0075539) was detectable only in the pupae and adults but was absent in larval stages. Out of the four isoforms however, DCP2_RA and DCP2_RE is observed to be expressed throughout development but DCP2_RE appears to be the most abundant and ubiquitous isoform expressed. Although, DCP2-RB is driven by the same promoter which drives DCP2_RA and RE, its absence does not necessarily indicate dearth of expression. *In-silico* analyses and data mining from the Eukaryotic Promoter Database (Dreos et al, 2014, 2017) indicate that DCP2-RD may be driven by the second promoter of DCP2 (DCP2_2). The protein isoforms coded by DCP2_RB and DCP2_RD are identical in sequence and size, but the exclusive expression of DCP2_RD in the larval gonadal tissues (ovaries and testes) and at a very low titre in the adults may be owing to the promoter being responsive to transcription factors in the gonadal tissues only and/or a putative undiscovered function of the transcript therein, despite absence of visible quantities of DCP2_RB.

**Figure 1:**
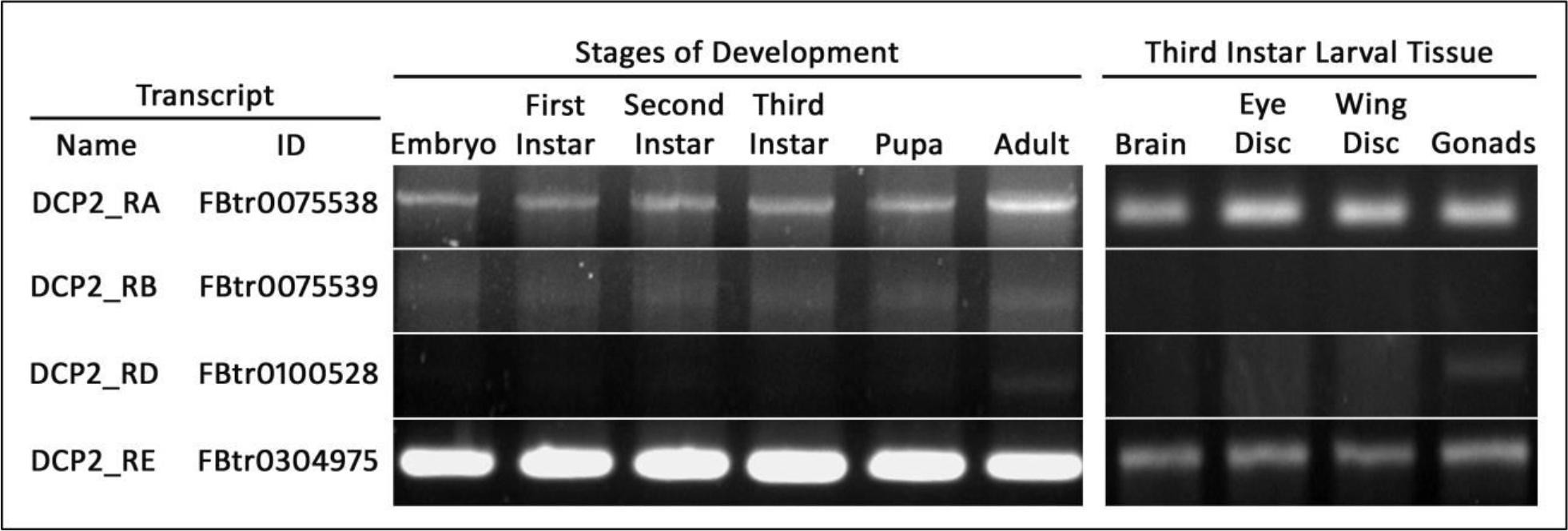
Electrophoretogram showing the expression pattern of different isoforms or splice variants of *DCP2* across *Drosophila* development and in selected third instar larval tissues.

### *DCP2* expresses in cells of diverse developmental lineages in the *Drosophila* embryo and the *DCP2* promoter *vis-à-vis DCP2* is active since early development

Evolution has been parsimonious in designing genes and ascribing roles to them and hence, determination of gene functions becomes incomplete without identification of the expression dynamics of the gene. In order to determine the endogenous expression pattern of *DCP2* in *Drosophila melanogaster*, we used the *DCP2*^*GAL4*^ (*DCP2*^*BG01766*^) which has a P{GT1}construct (Lukacsovich et al, 2001) bearing a GAL4, immediately downstream to the *DCP2* promoter. Using GFP as a reporter, we detected extremely robust signal in the late embryonic stages, wherein it expresses strongly in the embryonic epithelia (ectoderm) (**Fig. 2A**), the central nervous system (neuro-ectoderm) (**Fig. 2B**) and the dorsal muscles (mesoderm) (**Fig. 2C**) and is uniformly ubiquitous in all the segments in the embryo. With such a robust expression (of GFP), which is actually driven by the *DCP2* promoter, it is evident that *DCP2* is expressed and is active in embryonic cells derived from differing developmental lineages.

**Figure 2:**
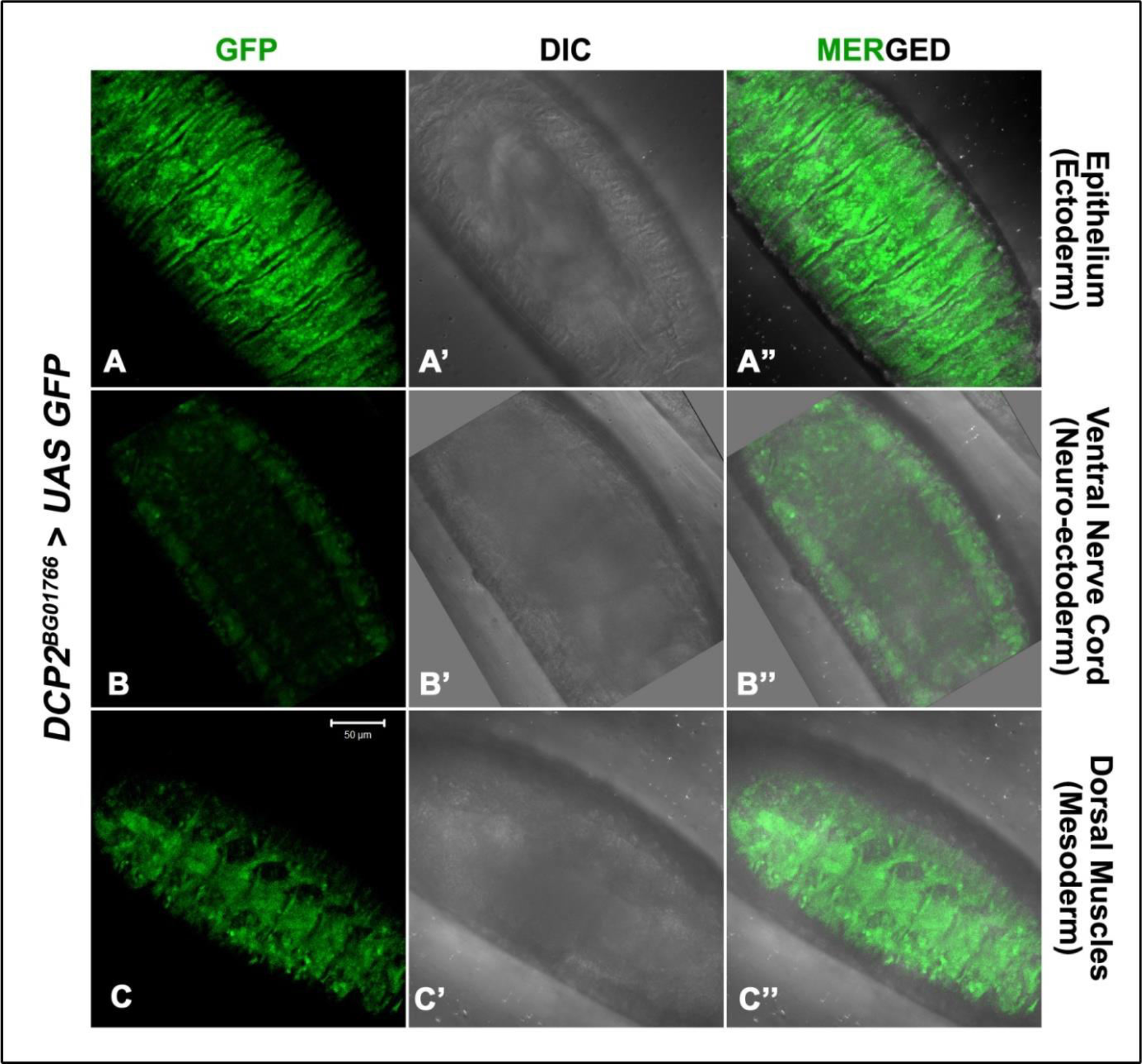
Confocal projections of late embryos (Stage 17) showing endogenous expression pattern of *DCP2* as determined by expression of GFP (green) by *DCP2*^*GAL4*^. Tissues of differing developmental lineages, *viz.*, ectoderm (A-A”), neuro-ectoderm (B-B”) and mesoderm (C-C”) show robust expression of GFP.

Dorsal closure is a major morphgenetic event during embryonic developemnt in *Drosophila* (Martinez-Arias, 1993) and involves an orchestrated interplay of numerous molecules (Lada et al, 2012) to drive the concerted movement of lateral epithelial cell sheets. With *DCP2* being expressed strongly in the embryonic epithelium, we investigated the possibility and nature of activity of the *DCP2* promoter during dorsal closure. To determine the real-time activity of the *DCP2* promoter during dorsal closure, we used a GAL4-responsive tripartite construct, G-TRACE (Evans et al, 2009). Using this transgenic line, in the embryonic stages (**Figure 3**), we observed that *DCP2* expresses in a more or less ubiquitous pattern very early in development, even prior to Stage 10 (**Figure 3A**). However, real-time expression was not detected in Stage 10, in which the germband is fully extended (**Figure 3B** and **3D**), and the Stage 12, wherein the germband is fully retracted (**Figure 3J** and **3L**). In the intervening stage, wherein the germband starts retracting, *i.e.*, Stage 11, we detect strong expression of *DCP2* (**Figure 3F** and **3H**). Again, when the epithelial sweeping initiates following germband retraction (Stage13), we see a surge in the RFP expression (**Figure 3N** and **3P**) which intensifies further in Stage 14, in which the lateral epithelia on either side are still moving (**Figure 3R and 3T**). This intense RFP expression is visible in Stage 16 as well (**Figure 3V and 3X**). The Stages 10 and 12 are developmental periods of low cell migration as against Stages 11, 13 and 14 wherein the epithelium moves as an initiative of collective cell migration and coordinated cell-shape changes. The expression potential of the *DCP2* promoter across DC revealed expression “crests and troughs”, such that the “crests” paralleled the periods in which cellular mobility or migration was maximal and *vice-versa* (**Figure 3Y**). The RFP activity is detectable only in stages which involve collective cell movement. The eGFP expression however depicts an early initial pulse of the gene expression which plausibly maintains a basal level of gene product. Hence, the dynamics of the promoter reflects a tightly regulated expression of *DCP2* and brings to light that *DCP2* may be an essential player during collective cell migration *vis-à-vis* epithelial morphogenesis.

**Figure 3:**
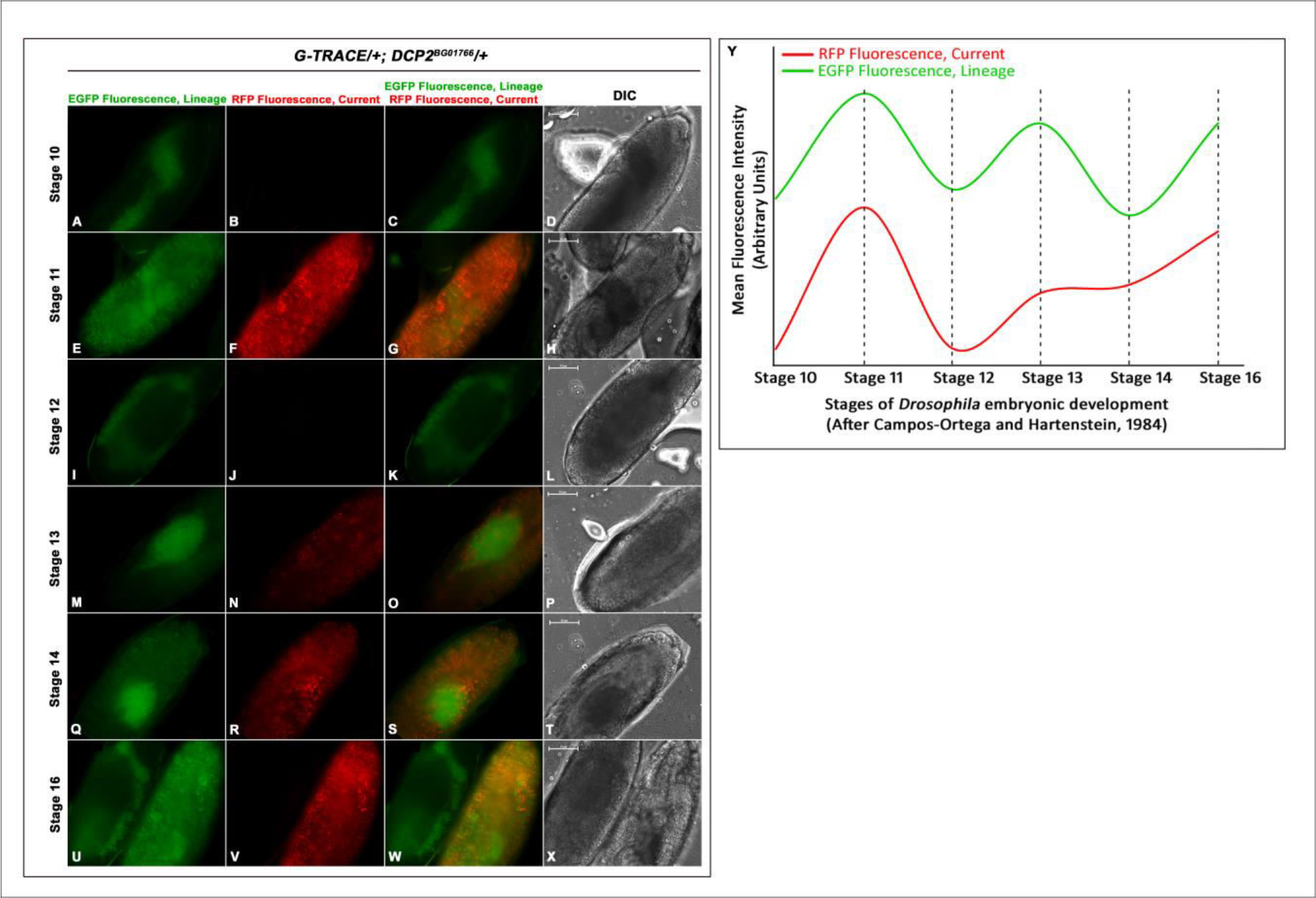
Lineage specific (EGFP) and real time (RFP) expression of *DCP2* in the embryonic stages using the GAL4-UAS based G-TRACE system. A-X show the expression pattern of the reporters along with the DIC images of the embryos. While real-time *DCP2* promoter activity is not detectable during Stages 10 and 12 and is low in Stage 13, it is robust in Stages 11 and 14. Y shows a plot of the fluorescence of both the reporters (GFP and RFP) across the stages observed, wherein a near-sinusoidal curve is obtained showing crests and troughs of *DCP2* promoter activity across the stages of Dorsal closure.

### DCP2 expresses in the amnioserosa and lateral epithelium during dorsal closure and its loss affects survival, epithelial morphogenesis and development of nervous system in the *Drosophila* embryo

Since *DCP2* shows active expression paradigms during embryonic dorsal closure (**Figure 9**), we examined the expression of DCP2 in the lateral epithelia in Stage 13 embryos along with the expression of activated JNK, a key mediator of dorsal closure (Jacinto et al, 2002), and Fasciclin III, a cell adhesion molecule (Bahri et al, 2010), both of which are expressed at the lateral epithelia and the leading edge (LE) cells. DCP2 was found to be expressed in the amnioserosa and throughout the lateral epithelium as well as in the cells at the LE (**Figure 4**). During the later stages of dorsal closure, parallel to the contra-lateral movement of the epithelia towards the dorsal side, the axon pathways are pioneered in the CNS across the ventral nerve cord, which form the complete nervous system by the end of Stage 16 (Bhuin and Roy, 2009). Examination of the ventral nerve cord also showed strong cytoplasmic expression of DCP2 (**Figure 5**).

**Figure 4:**
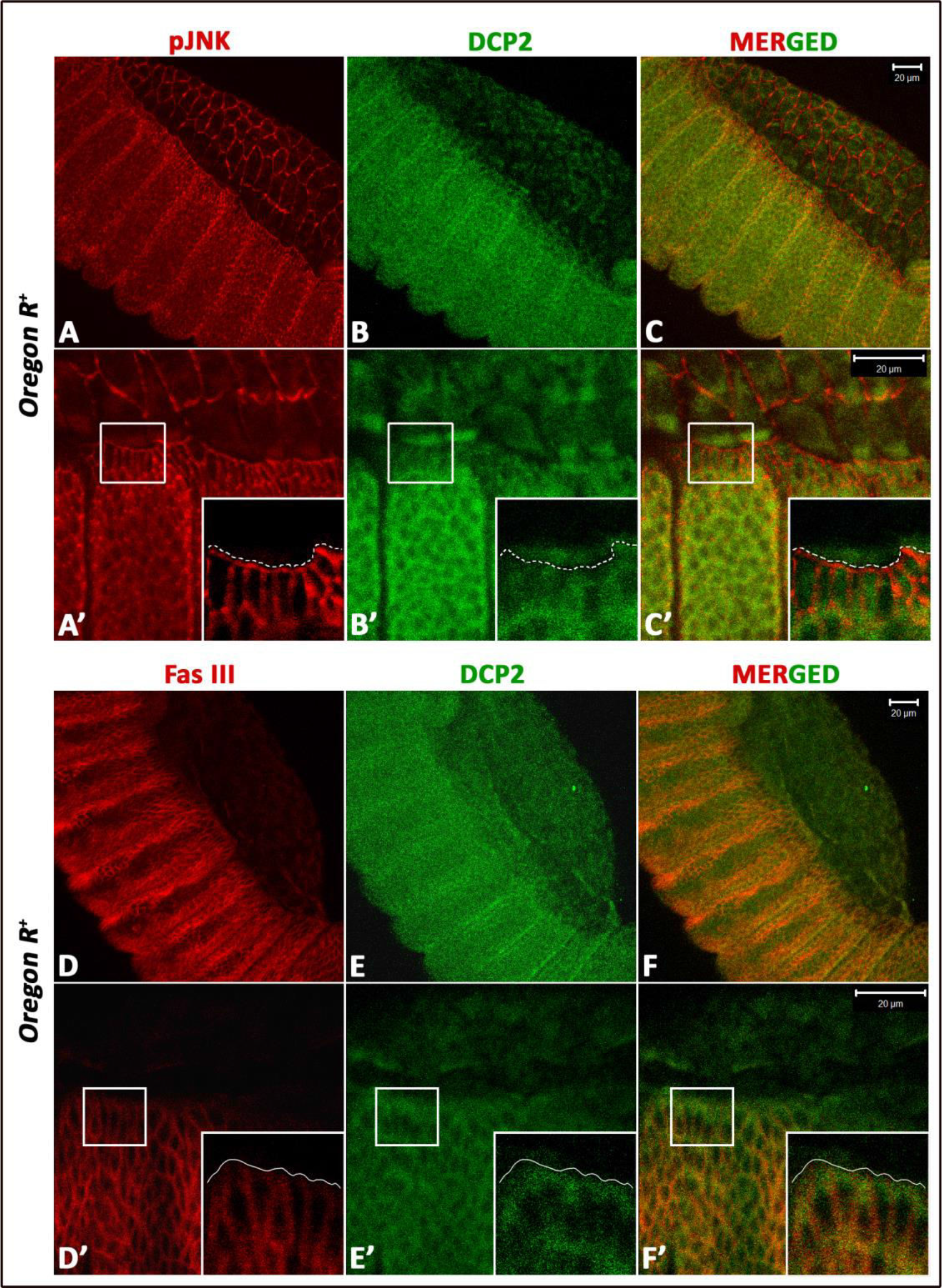
Confocal projections showing immunolocalisation of DCP2 in the amnioserosa and the lateral epithelium in Stage 13 embryos of wild type strain, co-stained for phospho-JNK (A-C) or the septate junction marker FasIII (D-F). In both cases, punctate expression of DCP2 in the lateral epithelium and amnioserosa (B and E) and at the leading edge (B’ and E’) is clearly visible.

**Figure 5:**
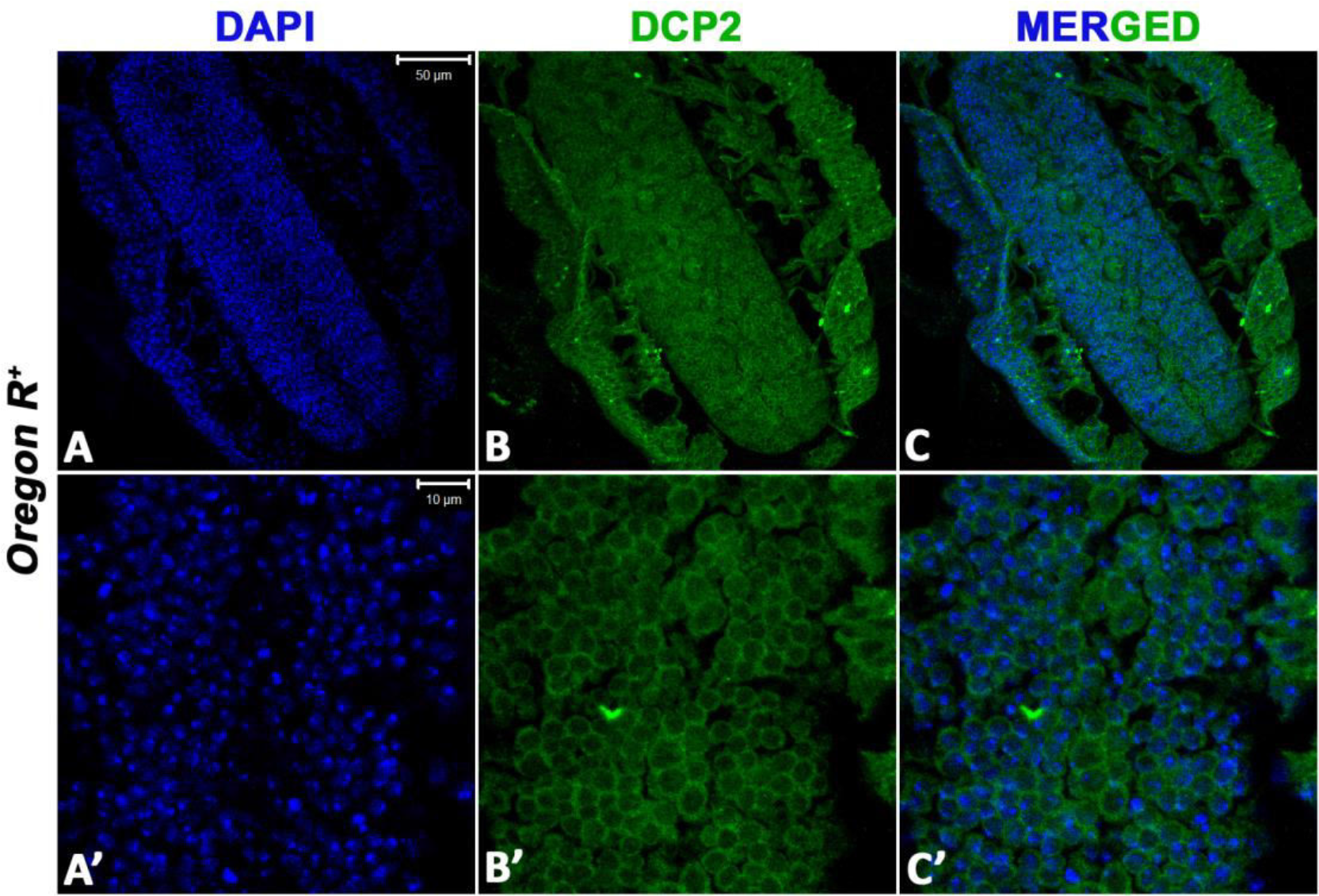
Confocal projections showing immunolocalisation of DCP2 in the ventral nerve cord of Stage 16 embryos of wild type strain show cytoplasmic expression of DCP2 (B and B’) in the ventral nerve cord. Nuclei are counterstained with DAPI.

We found that embryos homozygous for loss-of-function alleles of *DCP2* (*viz.*, *DCP2^BG01766^* and *DCP2*^*e00034*^) show embryonic lethality. *DCP2*^*BG01766*^ homozygotes are 100% embryonic lethal (N=500) whereas *DCP2*^*e00034*^ homozygotes show 12% lethality (N=500) at the embryonic stage and the remaining die before reaching the second instar larval stage.

### Defects in Epithelial Morphogenesis

Since we found strong expression of *DCP2* in the embryonic epithelium, we endeavored to explore whether a loss of *DCP2* function affects epithelial morphogenesis. Analysis of embryonic cuticles showed that all mutants displayed pronounced defects in epithelial morphogenesis patterns, ranging from defects in size, *viz.*, anterio-posterior or dorso-ventral dimensions, head involution defects and morphological defects, *viz.*, u-shaped or puckering (**Figures 6**). Since these defects are not mutually exclusive in that, a single mutant embryo could be displaying multiple defects at the same time, the morphological aberrations were scored individually first and then analysed for the presence of other concomitant defects. While 82.6% of the *DCP2*^*BG01766*^ homozygotes show altered anterio-posterior or dorso-ventral dimensions (*viz.*, elongated or compressed) of which 68.4% embryos are defective in head involution and 36.8% embryos are puckered, 21.7% embryos have gross defects in all the morphological parameters analysed. 4.3% of the embryos are exclusively puckered and 6.5% embryos show defects in head involution only. None of the *DCP2*^*e00034*^ homozygotes analysed was exclusively puckered. 80% of the embryos observed show altered dimensions out of which 12.5% are puckered and are defective in head involution and 62.5% embryos are not puckered but show head involution defects. 20% of the embryos observed show defects exclusively in dimensions or head involution. Figure 7 shows the above data represented with the help of a Venn diagram.

**Figure 6:**
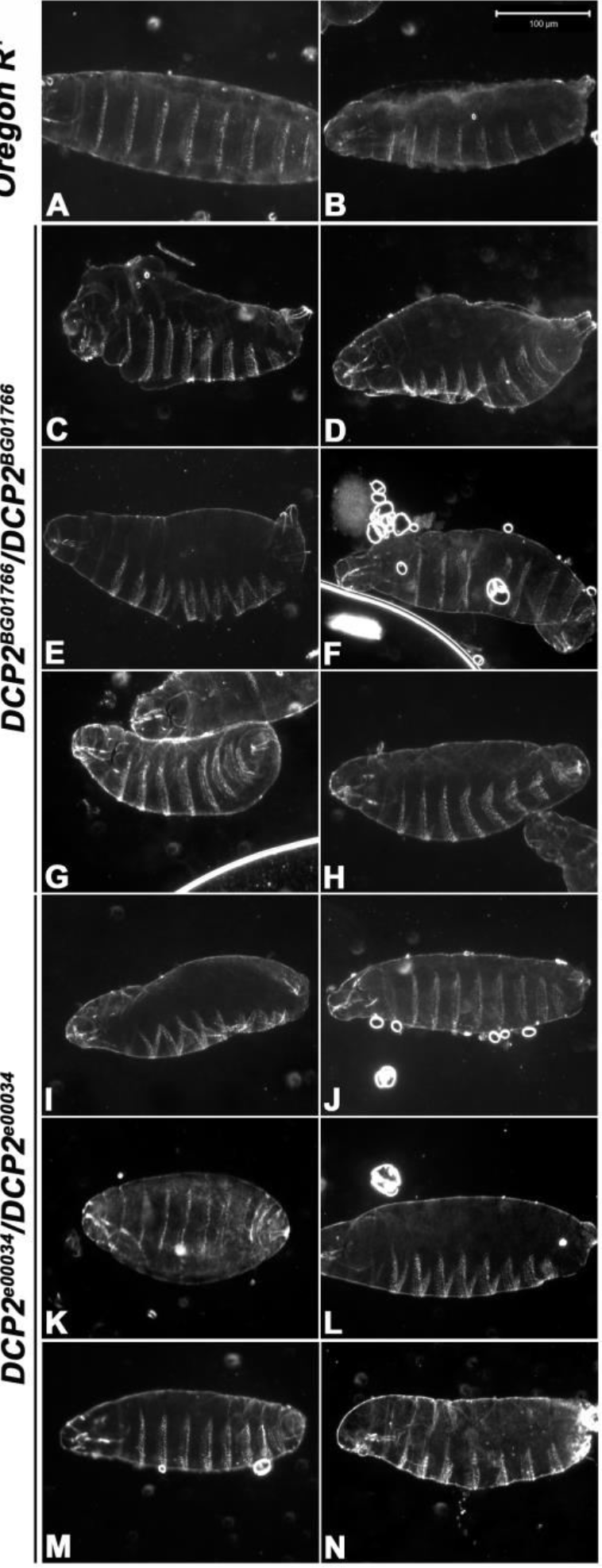
Dark field photomicrographs of embryonic cuticles of the wild type (A and B) and *DCP2* loss-of-function homozygotes, *viz.*, *DCP2^BG01766^* (C – H) and *DCP2^e00034^* (I – N). Note the altered dimensions and /or morphology and defects in head involution exhibited by the mutants as compared to the wild type.

**Figure 7:**
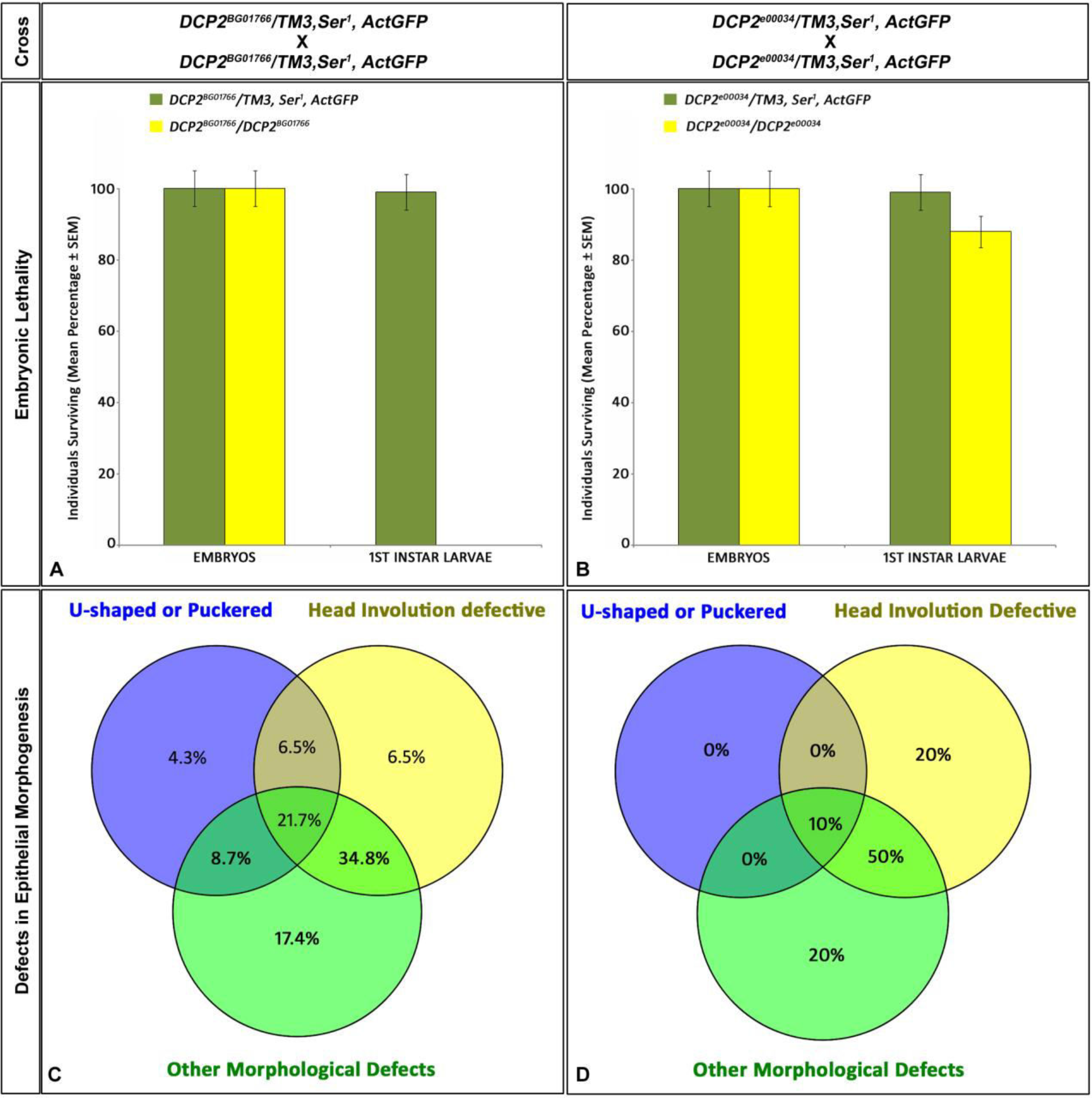
Embryonic Lethality and Defects in Epithelial Morphogenesis in *DCP2* loss-of-function homozygotes. *DCP2*^*BG01766*^ homozygotes are 100% embryonic lethal (A) and exhibit a broader range of epithelial morphogenesis defects being altered in anterio-posterior or dorso-ventral dimensions along with puckering and defective head involution (C), but *DCP2*^*e00034*^ homozygotes show only 12% lethality at the embryonic stage and display a milder range of defects with none of them being exclusively u-shaped or puckered (B).

### Defects in Nervous System development

We used mAbBP102, an antibody to mark all CNS axons (Seeger et al. 1993) such that the gross morphology of CNS in an embryo is revealed. In wild-type embryos, axons form an orthogonal structure having longitudinal axon tracts. These axon tracts run anterio-posteriorly, being positioned at either side of the midline, and a pair of commissural tracts joins the longitudinal pathways in each segment of the embryo (Bhuin and Roy, 2009). *DCP2*^*BG01766*^ homozygotes showed thinning of longitudinal connectives and compressed segmental commissures (**Figure 8 B and B’**) similar to the *karussell* phenotype (Hummel et al, 1999), whereas, *DCP2*^*e00034*^ homozygotes showed thinning of longitudinal connectives and lateral commissures (**Figure 8 C and C’**).

**Figure 8:**
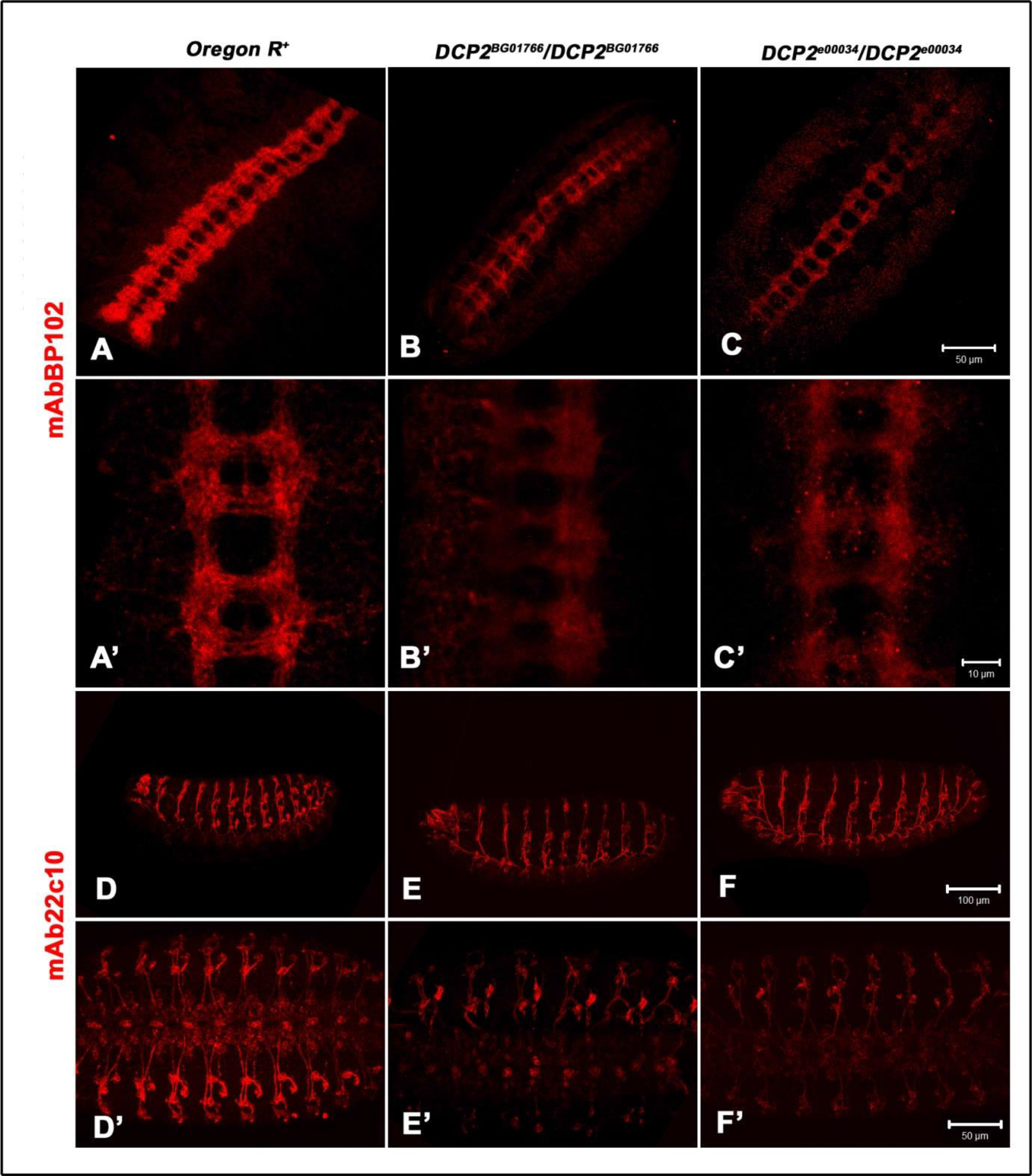
*DCP2* null homozygotes display defects in CNS and PNS organization. Upper panel: Wild type embryos, stained with mAbBP102 show regular arrangement of longitudinal connectives and segmental commissures (A and A’). *DCP2*^*BG01766*^ homozygotes showed thinning of longitudinal connectives and compressed segmental commissures (B and B’), whereas *DCP2*^*e00034*^ homozygotes showed thinning of longitudinal connectives and lateral commissures (C and C’). Lower panel: In the PNS, axons run from the ventral nerve cord to the periphery of the embryos in each hemisegment (D and D’). A loss of *DCP2* causes misrouting and collapse of fasciculating axons (E and E’; F and F’).

In order to study the embryonic PNS axons further, mutant embryos were stained with mAb22C10, which recognizes the microtubule-associated protein, futsch (Hummel et al. 2000). It labels all the cell bodies, dendrites, and axons of all PNS neurons, and a subset of neurons in the VNC of the CNS (Fujita et al. 1982). Therefore, defects, such as the disruption of the nervous system, the collapse of the axon tracts, fasciculation defects/thinning of axons, and the loss or gain of neurons can often be distinguished. In the wild type embryos, each segment contains three highly stereotyped clusters of PNS neurons connected by axon bundles. In the mutants, misrouting of axons and collapsed axons could be detected (**Fig. 8 E, E’ and F, F’**), which were absent in the wild type, implying a role for DCP2 in the fasciculating axons.

### *DCP2* loss-of-function mutants show elevation and spatial perturbation of JNK signaling

The JNK signaling cascade is an essential player of *Drosophila* gastrulation *vis-à-vis* embryonic development wherein it modulates important events such as dorsal closure (Jacinto et al, 2002; Kushnir et al., 2017) and architecture of the nervous systems (Shklover et al, 2015; Karkali et al, 2016). Since *DCP2 loss-of-function* mutants show defects associated with either process, we wanted to investigate the spatial expression of the JNK cascade. We harnessed the bio-sensor, TRE-RFP to identify the spatial pattern of JNK signaling (Chatterjee and Bohmann, 2012) in the wild type embryos and in embryos homozygous for *loss-of-function* alleles of *DCP2*. Both *DCP2*^*BG01766*^ and *DCP2^e00034^* homozygotes showed enhanced RFP expression, implying an elevation in the JNK signaling cascade. Further, the pattern of RFP *vis-à-vis* JNK signaling is spatially disrupted in the *DCP2* mutants, implying a perturbation and/or misregulated JNK signaling (**Figure 9**). In the wild type embryos it is expressed at the juncture of the two epithelial sheets following completion of dorsal closure, mimicking the stitch at a suture, whereas the mutant homozygotes show a spatial distortion of JNK signaling.

**Figure 9:**
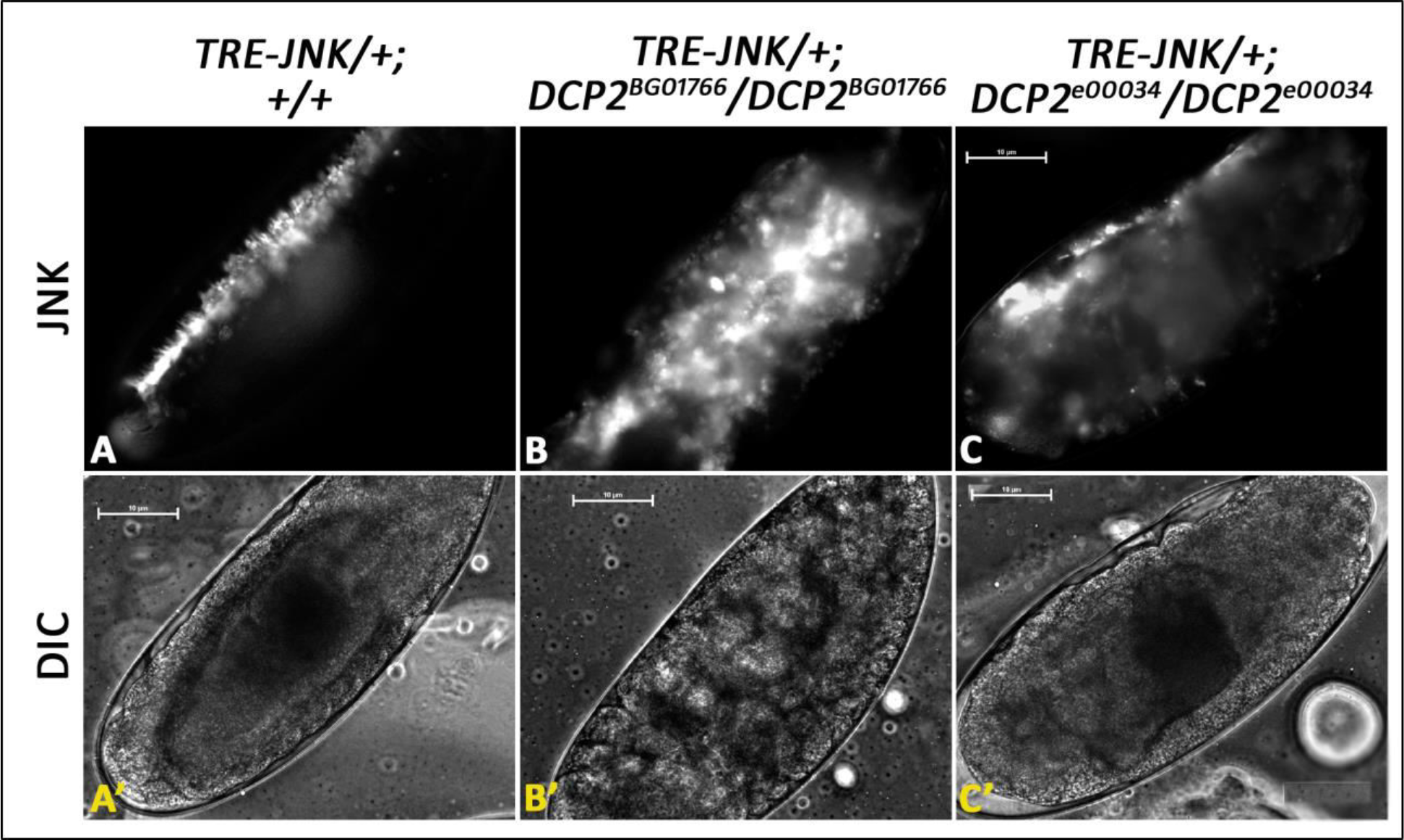
Expression of JNK as determined by the biosensor, TRE-JNK in Stage 15 embryos of wild type and *DCP2* loss-of-function mutants. While JNK appears as a suture in the wild type embryos (A and A’), it’s spatial expression is completely disrupted in *DCP2* mutant homozygotes (B and C).

In the developing *Drosophila* embryo undergoing gastrulation, epithelial morphogenesis and axonogenesis are morphogenetic events of utmost importance that require a well orchestrated spatio-temporal regulation of gene expression. During initiation of dorsal closure wherein the two lateral epithelia initiate contra-lateral movement to eventually seal the dorsal opening, the dorsal-most lateral epithelial cells express high levels of JNK (Noselli and Agnes, 1999). The JNK signaling pathway is a core signaling pathway in the process of dorsal closure at the time of gastrulation in *Drosophila* embryos (Noselli, 1998; Noselli and Agnes, 1999; Ramet et al 2002; Stronach and Perimon, 2002). While DCP2 co-expresses with JNK ubiquitously on the dorso-lateral epithelia, the leading edge and the amnioserosa, monitoring the activity of the *DCP2* promoter in real time across the stages of dorsal closure shows spurts of promoter firing in the stages which involve large scale cell migration and movement. Such subtle and precisely timed gene activity is indicative of thorough fine-tuning of the expression of *DCP2*. The ablation of *DCP2* does not lead to “dorsal open” embryos, but generates a spectrum of defects including altered embryonic dimensions and defects in head involution, improper fasciculation of axons and defects in segmental commissures and longitudinal connectives, causes embryonic and larval lethality implies significant perturbation in these developmental gene expression programs and indicate a more concerted and fundamental role of *DCP2* in regulating these phenomena. It is worth mentioning that despite ubiquitous expression of *DCP2*, the ectodermal (epithelium) and neuro-ectodermal (CNS and PNS) tissues are most affected following ablation. It is interesting to note that while the nematode worm is a closer relative of the fly in the evolutionary tree, the embryonic lethality and the developmental defects are similar to those observed in a distant relative, *Arabidopsis* (Xu et al, 2006; Goeres et al, 2007). Since the JNK signaling pathway is fundamental to the process of dorsal closure during gastrulation in *Drosophila* embryos (Noselli, 1998; Noselli and Agnes, 1999; Ramet et al, 2002; Stronach and Perrimon, 2002) and both epithelial morphogenesis and CNS development are dependent on JNK activity (Jacinto et al, 2002; Kushnir et al., 2017; Shklover et al, 2015; Karkali et al, 2016), the altered expression patterns of JNK or misdirected JNK signaling under the influence of loss of *DCP2* in the different alleles could be a probable cause of the defects observed in each case.

### The *DCP2* promoter shows consistent developmental activity in the larval tissues along with tissue-specific expression paradigms of the translated protein

Real-time activity of the promoter in the various larval tissues, with a better insight, demonstrates that despite expression since early development, the *DCP2* promoter is active during late stages of larval development as well. In the larval tissues (118±1 h ALH), we could identify a consistent GFP expression in the larval brain, eye-antennal disc, salivary gland and wing discs. Besides prior developmental activation and ubiquitous expression, the *DCP2* promoter shows enhanced activity/expression in specific cells in the brain (**Figure 10 B and C**) and eye disc (**Figure 10 G and H**). Although, the ventral ganglion depicts an overlap of prior and real-time activity (**Figure 10 B-D**), the cerebral hemispheres show selective expression in real-time, which is limited to cells of the antennal lobe and the Kenyon cells (**Figure 10 B’-D’**). Similarly, the cells in constituting the antennal disc show a more consistent *DCP2* activity, exemplified the greater degree of overlap of the reporters (**Figure 10 G-I**). However, the cells of the eye disc show heterogeneity of reporter expression, *viz., DCP2* is active in all the ommatidia but certain cells show a transient activity at the stage observed (**Figure 10 G’-I’**). While the salivary gland nuclei show a complete colocalization of reporters with similar intensity (**Figure 10 L-N**), wing discs show a lower expression of eGFP as compared to RFP.

**Figure 10:**
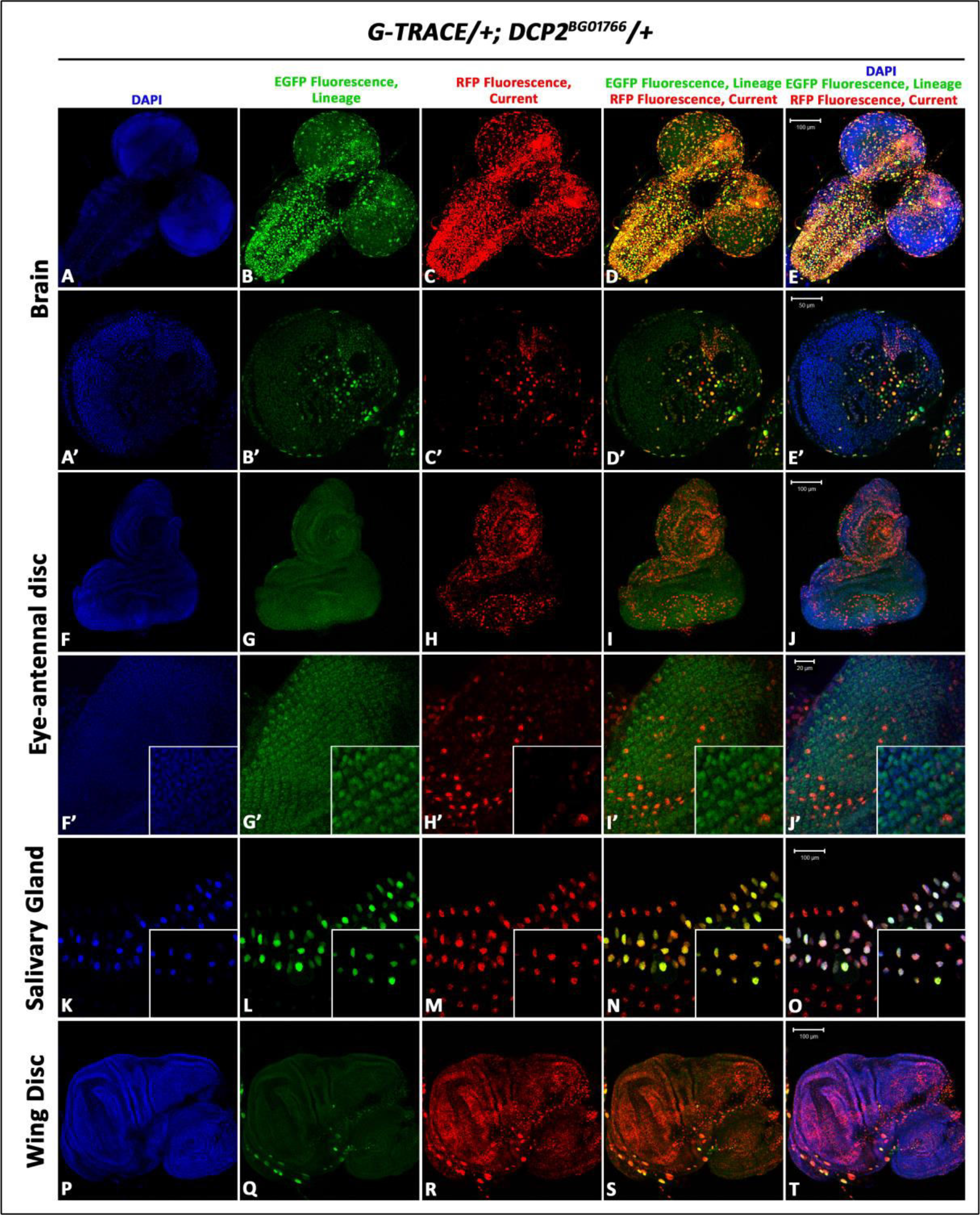
Lineage specific (EGFP) and real time (RFP) expression of *DCP2* in the larval tissues using the GAL4-UAS based G-TRACE system. Although the ventral ganglion (B and C) and the antennal disc (G and H) show significant overlap of the reporters, the central brain (B’ and C’) and the eye-disc (G’ and H’) show heterogeneity of expression. The salivary gland nuclei and the wing disc show strong real-time expression of the *DCP2* promoter along with prior developmental expression.

Since, the GAL4 is driven by the *de novo* promoter of *DCP2*, which in absence of the GAL4 coding region, would have transcribed the gene *per se*, this transgenic construct, *viz.*, G-TRACE allows the spatio-temporal expression potential of the promoter to be determined. Thus, the expression of the reporters (GFP and RFP) may be directly correlated with the gene expression pattern or potential in the wild type individual and hence, *via* the GAL4, directly demonstrates the spatio-temporal gene expression dynamics.

### Brain

In the larval brain, immunolocalisation of DCP2 shows a uniform cytoplasmic expression throughout the dorso-ventral and anterio-posterior axes of the tissue (**Figure 11A’, B’ and D’**). However, significantly high levels were detected in a subset of neurons in the ventral ganglion (**Figure 11A’ and D’**) and in a cluster of neurons in the dorso-lateral and dorso-medial region of the central brain (**Figure 11A’ and C’**), which are proximal to the Mushroom Body as well as in the Kenyon cells (**Figure 11K and O**). However, it is completely absent from the most prominent neuronal structures *viz.*, the Mushroom Body in the central brain and in the neurons of the optic lobe (**Figure 11M-O**).

**Figure 11:**
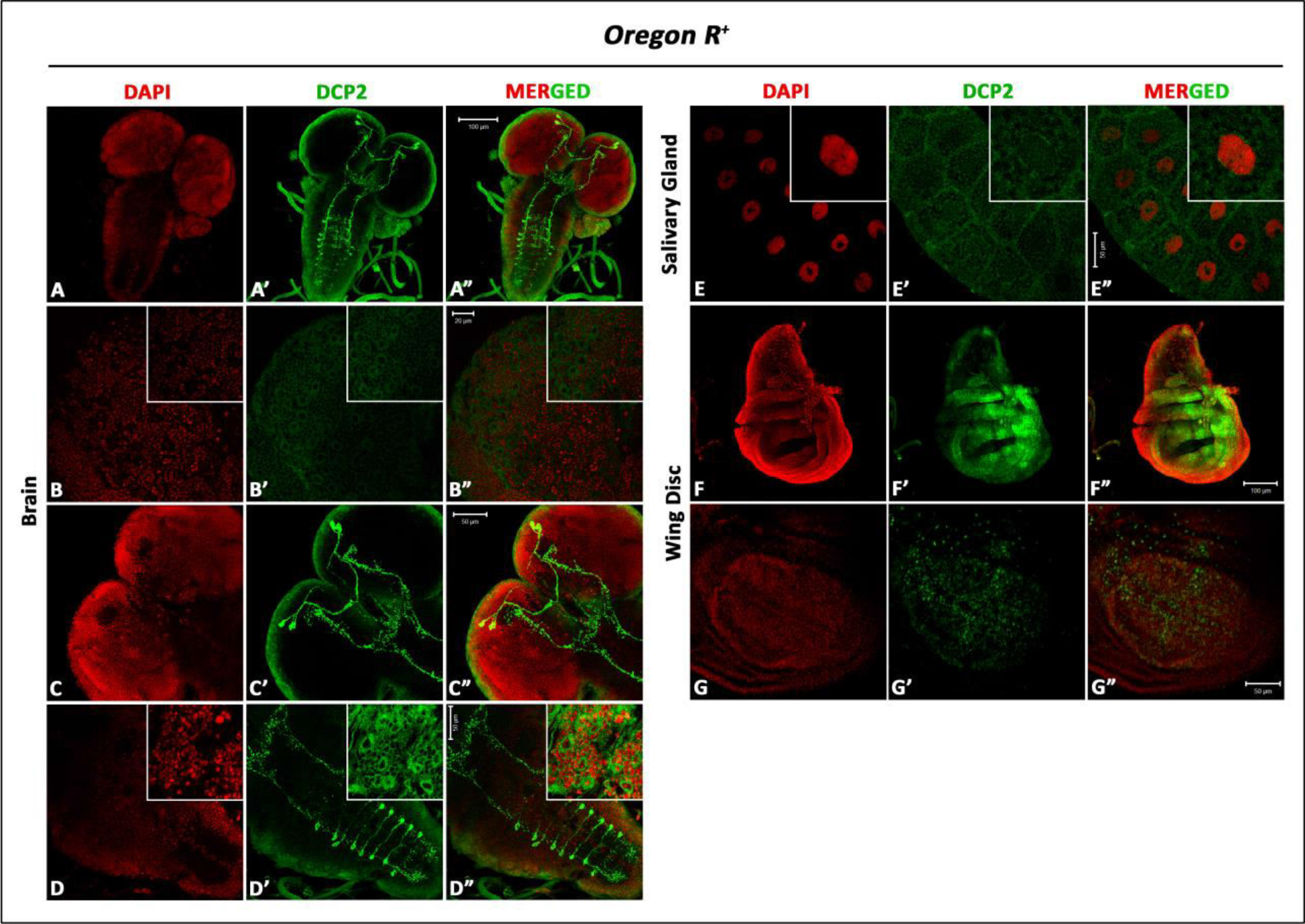
Confocal projections showing immunolocalisation of DCP2 in the larval tissues. A-A” shows the expression pattern in the larval brain. B, C and D show higher magnifications of the same, wherein we find a ubiquitous cytoplasmic expression of DCP2. Visible in C’ and D’ are a subset of neurons which show high expression of DCP2. In the salivary glands (E), besides cytoplasmic expression, the vesicles in the cytoplasm appear to be arduously decorated with punctate distribution of DCP2. F shows the pattern of expression of DCP2 in the wing disc. Shown in G is a confocal section which shows a magnified view of the wing pouch wherein DCP2 at the anterio-posterior and dorso-ventral margins in a cruciform pattern.

### Salivary Glands

In the salivary glands, DCP2 shows a punctuate distribution in the cytoplasm and decorates the nuclear and cellular membranes arduously (**Figure 11E’ and E”**). The cytoplasmic vesicles appear bounded by bodies rich in DCP2 (**Figure 11E”**). Since DCP2 is a cognate resident of the P-bodies, it may be fair enough to interpret the cytoplasmic network of DCP2 punctae in the glands as the pattern of P-bodies which are essential for maintaining transcript homoeostasis.

### Wing discs

The wing discs show very strong expression of DCP2 in the pouch region as compared with the notum (**Figure 11F’**), besides the uniform ubiquitous cytoplasmic distribution similar to that observed in other tissues. Most notably, the expression of DCP2 in the central sections of the pouch overlaps with the expression of the Anterio-Posterior determinant Decapentaplegic (Dpp) (Zecca et al, 1995) and the Dorso-Ventral determinant Wingless (Wg) (Neumann and Cohen, 1997), thereby presenting a “cruciform” pattern in the pouch (**Figure 11G’**), which may be essential during the morphogenesis of the wing blade.

Immunolocalisation of DCP2 to the cytoplasm in all the tissues examined across development recapitulates the results observed in similar studies in the nematode worm, *C. elegans* (Lall et al, 2005) and in the thale cress, *Arabidopsis* (Xu et al, 2006). In spite of uniform ubiquitous cytoplasmic expression in the larval tissues, certain paradigms of expression have been noticed. The protein shows distinct punctate expression of the protein in the wing imaginal disc along the anterio-posterior and dorso-ventral axes in the wing pouch, mimics the expression patterns of the TGF-beta homologue Decapentaplegic and Wingless, respectively. In the salivary glands as well, the protein is cytoplasmic but shows high titres at the membranes. Being the sole decapping agent in *Drosophila*, DCP2 is expressed ubiquitously throughout development but, the selectively high expression in certain cell types in the brain or the wing pouch point towards some yet unknown “moonlighting” functions of *DCP2* in the development and maintenance of cellular homoeostasis in these tissues.

### DCP2 shows high expression in the Corazonin neurons in the larval CNS

Besides ubiquitous expression, DCP2 has a typical expression paradigm in a subset of neurons in the larval CNS. In order to identify/type the DCP2 immunopositive neuron(s) in the larval ventral nerve cord (VNC), we tried mapping them against the Fasciclin II (FasII) landmark system (Santos et al, 2007) (**Figure 12**). Comparing the FasII “coordinates” with the DCP2 expression paradigm, we observed that DCP2 expresses in a cluster of three neurons in the Dorso-lateral (DL) region and in a neuron located medial to the DL neurons (Dorso-medial; DM) in the central brain, and in eight pairs of bilateral neurons in the ventral nerve cord. The neurons in the ventral ganglion correspond to a subset of the thoracic (T2 and T3) and abdominal (A1 – A6) neuromeres. Although DCP2 is absent from the most prominent neuronal structures expressing FasII, *viz.*, the Mushroom Body and the neurons innervating the eye, in the central brain (**Figure 12 D-F**) and in the Dorso-lateral and Dorso-medial longitudinal tracts in the VNC (**Figure 12 G-I**), the DL neurons appear to innervate the Ring gland and the aorta. Lateral views of the central brain (**Figure 12 D’-F’**) and the VNC (**Figure 12 G’-I’**) show that the DCP2-positive neurons lie below the Fas II immunopositive Dorso-lateral and Dorso-medial longitudinal tracts but ascend above the Mushroom Body (MB) in the central brain.

**Figure 12:**
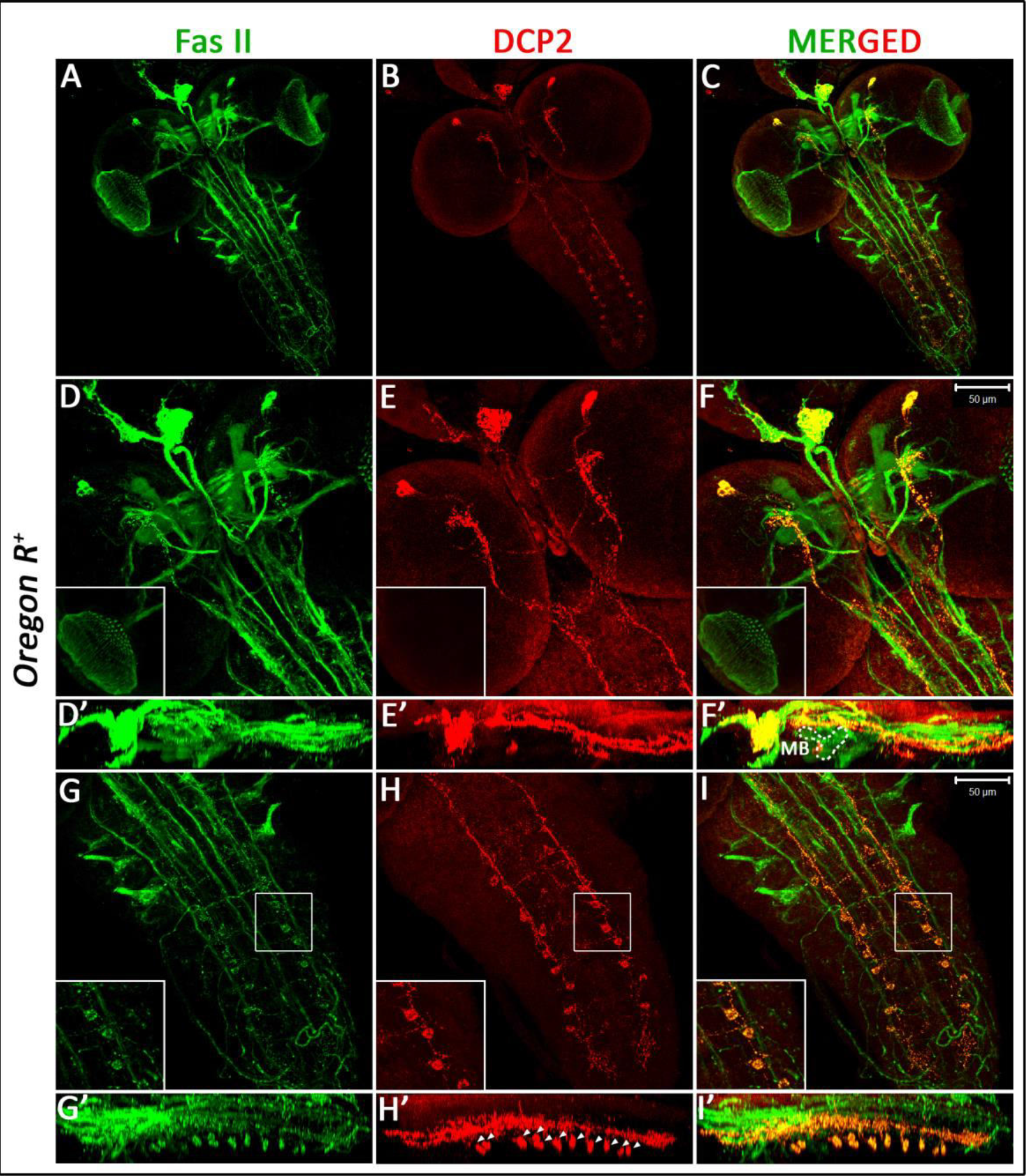
Mapping of the neuron(s) expressing high titres of DCP2 in the whole mount preparations of the larval brain (A-C) in the FasII landmark system (Santos et al, 2007). Note the absence of DCP2 in the neurons of the optic lobe (inset D-F). Z-axis stacks show that the DCP2 positive immunopositive neuronal tracts lie below the FasII immunopositive tracts in the larval ventral ganglion but ascend over the Mushroom Body (MB) in the central brain (D’-F’). However, a subset of thoracic and abdominal neuromeres co-express FasII and DCP2 (G-I). Z-axis stacks (G’ – I’) showing lateral view of the larval ventral ganglion depicted in G-I demonstrate co-expression of FasII and DCP2 in the neuromeres.

While DCP2 did not show co-expression with the different neuropeptides *viz.* Crustacean Cardioactive Peptide (CCAP; Veverytsa and Allan, 2012), Drosophila Insulin like Peptide-2 (Dilp2; Liu et al, 2016), Corazonin (Crz; Lee et al, 2008) and short Neuropeptide F (sNPF; Nassel et al, 2008), biogenic amines Tryptophan hydroxylase (TH; Friggi-Grelin et al, 2003) and Dopamine decarboxylase (Ddc; Vomel and Wegener, 2008), or the transcription factor Apterous (Ap; Rincon-Limas et al, 1999) (**Supplementary Figure 2**), complete colocalization or overlap of expression was observed with the neurons expressing Corazonin (**Figure 13**), while only the Dorso-lateral neurons showed co-expression with sNPF (**Supplementary Figure 3**). Corazonin neurons constitute three neuronal subsets, *viz*., the dorso-lateral (DL) and dorso-medial Crz neurons (DM), and the Crz neurons in the ventral nerve cord (vCrz) (Lee et al, 2008), are essential for combating stress (Zhao et al, 2010) and co-express the transcription factor Apontic, which is necessary and sufficient to mediate sensitivity to ethanol (McClure, 2013).

**Figure 13:**
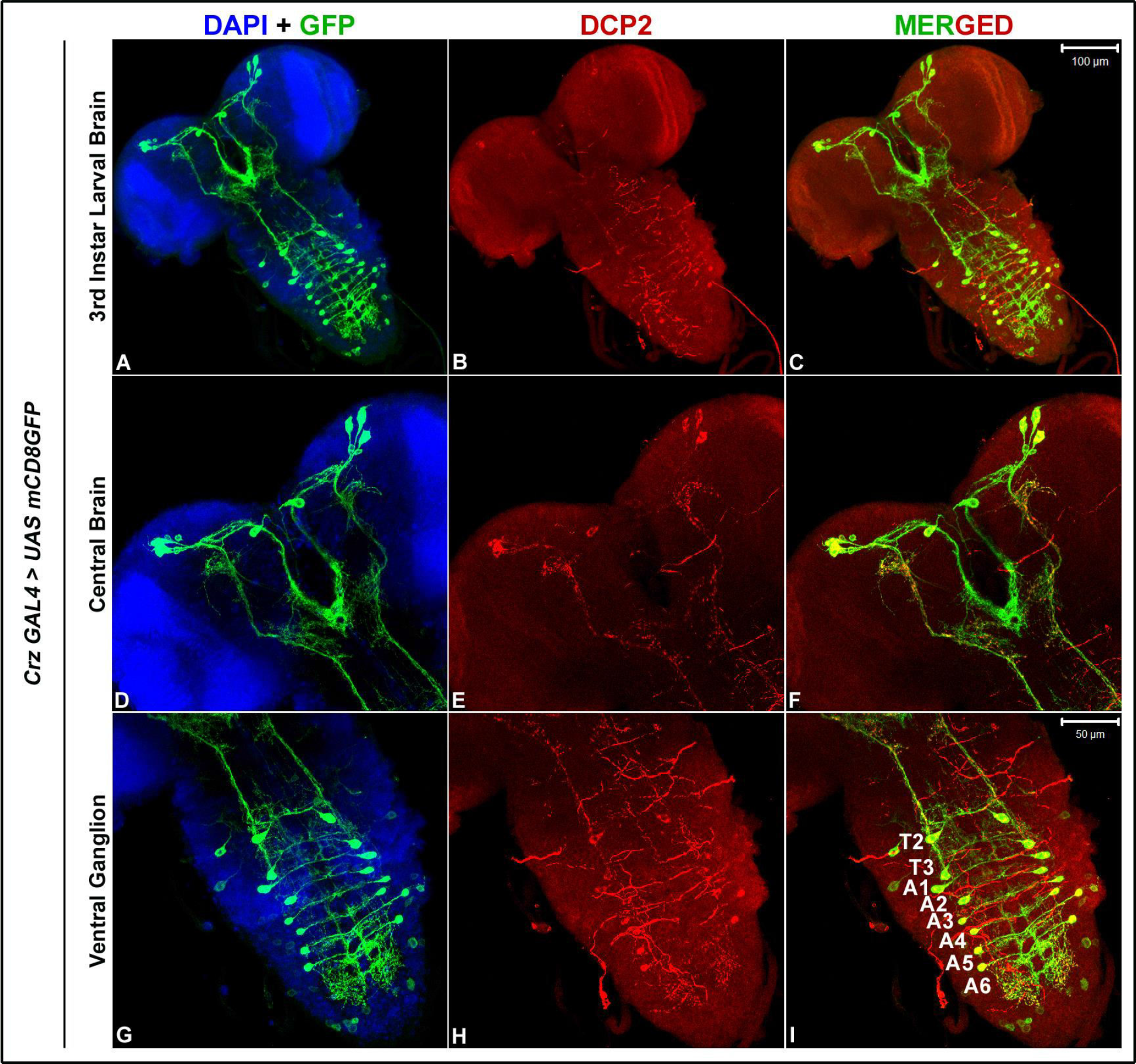
Mapping of the neuron(s) expressing DCP2 in the whole mount preparations of the larval brain against the Corazonin expressing neurons. A-C shows the expression pattern of DCP2 and Corazonin neurons in the larval brain. D-F and G-I show magnified view of the central brain and the ventral ganglion respectively, where complete colocalization of DCP2 and Corazonin is visible.

### Knockdown of *DCP2* in the Corazonin neurons reduces sensitivity to Ethanol

We further asked whether *DCP2* function in the Corazonin neurons is required for their activity. When *DCP2* was knocked down in these neurons specifically, it did not affect the morphology, pathfinding or architecture of the Crz neurons in the larval brain (**Supplementary Figure 4**) but, delayed and/or reduced the sedation sensitivity to ethanol in the adult flies. The time to 50% sedation (ST50) was calculated to be ~8.5 min for the control flies (N=200), while the *DCP2* knocked down flies showed an ST50 of ~13.5 min (N=200). While the control flies are sedated completely in ~12 min, the *DCP2* hypomorphs are active till ~19 min (**Figure 14 A and B**). This demonstrates the reduced sensitivity of the *Crz GAL4>UAS DCP2* RNAi knocked down flies to ethanol.

**Figure 14:**
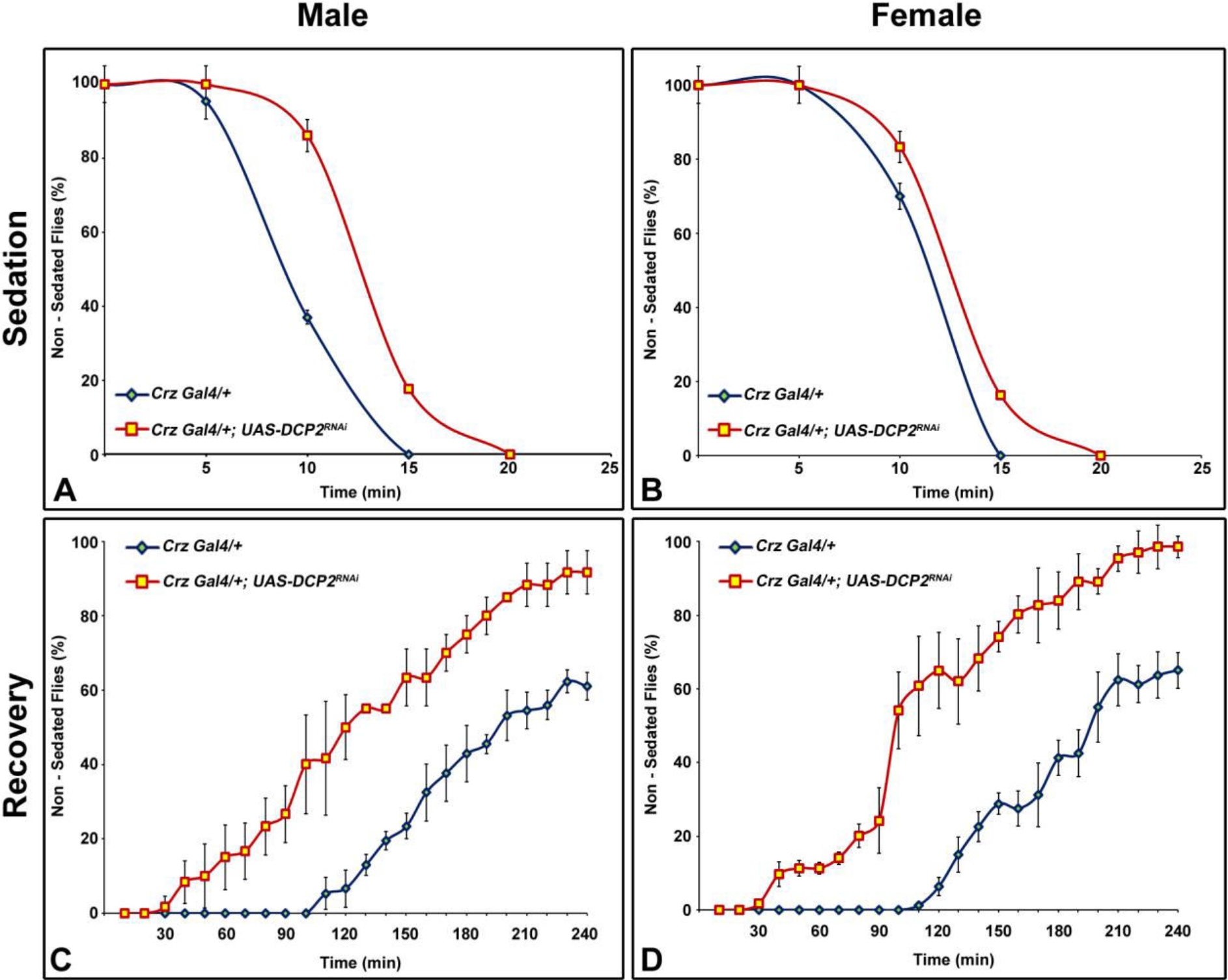
Graphs showing the response to Ethanol induced sedation (A and B) in the control (*Crz-Gal4*/+; blue lines) or *DCP2* knocked down (*Crz-Gal4*/+; *UAS-DCP2*^*RNAi*^/+) flies (red lines) and recovery from the same (C and D). The knocked-down flies (both male and female) show reduced sensitivity to ethanol vapours as is visible from their delayed sedation behaviour (A and B) or enhanced recovery from sedation (C and D).

Following sedation, the DCP2 hypomorphs showed early onset of recovery (~30 min) as against the control flies, which started showing activity/onset of recovery after ~110 min. While the control flies started assuming normal standing posture in ~2h, the *DCP2* knocked down flies showed significant early recovery, with ~80% of the flies recovering by 3h as against ~40% recovery exhibited by the control flies in the same time (**Figure 14 C and D**). Also, sedation induced death was higher in the control group as compared to the *DCP2* hypomorphs. During sedation and recovery phases, no sex specific differences were observed in flies of either genotype. These results suggest that *DCP2* function is required in the Crz neurons for regulation of ethanol related behaviour and ethanol metabolism.

The Crz neurons require the transcription factor Dimmed and its target enzyme, Peptidylglycine-alpha-hydroxylating monooxygenase (PHM) (Park et al, 2008) for synthesis of Corazonin while the transcription factor Apontic (Apt) is necessary and sufficient for regulating the activity of the Crz neurons and/or release of Corazonin during ethanol exposure (McClure, 2013). The delay in sedatory behaviour during ethanol exposure and the quick recovery from sedation demonstrate impaired function of the Crz neurons and/or perturbed corazonin signaling following knockdown of *DCP2* in the Crz neurons, although the mechanism behind such altered physiology remains unknown.

## Summary and Conclusion

Analysis and identification of the expression patterns of genes and/or proteins in model organisms across the evolutionary tree are important for understanding the spectral paradigm of gene function. The extent to which a gene and its expressome are conserved across diverse organisms indicates the precision of its function across phyla. mRNA decapping proteins are present in all metazoans and serve to initiate the decay of mRNA and are therefore important for regulation of gene expression *vis-à-vis* cellular physiology. The patterns of expression and paradigm range of physiological aberrations following the ablation or knockdown of *DCP2* is indicative of the fundamental regulatory role played by it during development and bring to light the hitherto undiscovered plausible novel functions of DCP2. It is yet unknown as to whether it’s function in the modulation of developmental events is *via* the *de novo* function of mRNA decapping or is a manifestation of moonlighting behaviour (Mani et al, 2014). Summarizing the present observations, our findings demonstrate that *DCP2* plays a major modulatory function in developmental gene expression and is essential for maintenance of organismal physiology at all stages of development.

## Acknowledgements

The authors acknowledge the fly community for generously providing fly stocks. We thank Prof. B. J. Rao, TIFR, Mumbai for providing the *TRE-JNK/CyO* stock, Prof. Gaiti Hasan, NCBS, Bangalore for providing the *sNPF-GAL4*, *Dilp2-GAL4* and *Crz-GAL4/CyO* stocks and Prof. Utpal Banerjee, UCLA for providing the *G-TRACE/CyO* flies. We duly acknowledge the National Facility for Laser Scanning Confocal Microscopy, Department of Zoology, Banaras Hindu University. Financial support from DST-FIST, UGC-UPE and CAS Zoology are duly acknowledged. We sincerely acknowledge Nabarun Nandy for assistance, valuable discussions and proofreading the manuscript. We sincerely thank Department of Science and Technology (DST) for providing INSPIRE Fellowship to RK.

## Author Contributions

RK, conceptualization, resources, methodology, investigation, data curation, formal analysis and interpretation, writing the manuscript. JKR, supervision, resources, analysis and interpretation, writing the manuscript.

## Conflict of Interest

The authors declare no conflict of interest.

