## Supplemental Information for "*DCP2* plays multiple roles during *Drosophila* development – possible case of moonlighting?"

**Supplementary Information**

| PRIMER | SEQUENCE (5′ → 3′) | Amplicon Size (in bp) | | | | |
|---|---|---|---|---|---|---|
| | | cDNA | | | | gDNA |
| | | RD | RB | RA | RE | |
| D_F<br>DBAE_R | ACAACGATTCAATACATATACAGCT<br>CTGTTTTTGTTGCTCGTGTTGT | 165 | – | – | – | 343 |
| BAE_F<br>DBAE_R | GCAATTTAGATCGCGAAAAAGTTC<br>CTGTTTTTGTTGCTCGTGTTGT | – | 159 | 974 | 974 | 2455 |
| EU_F<br>EU_R | TCATTTGTCTGGGCCAAGTGAC<br>TGGGATTGCAGTTCATCAAATG | – | – | – | 233 | 233 |
| EXON_F<br>EXON_R | TCAGTACT*GGTACC*GACTACATCGAGGCGTTCATCA<br>TCAGTACT*GGTACC*GTCATTGGCGTTGCTATTGTAGT | 618 | 618 | 618 | 618 | 618 |

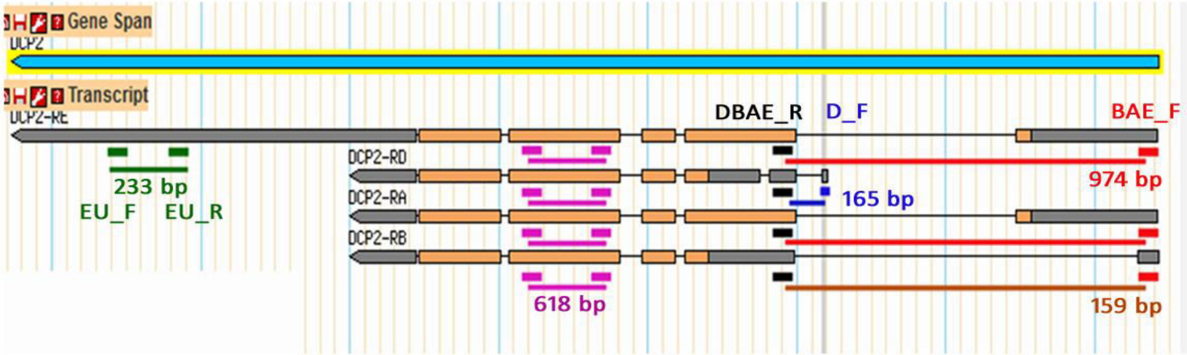


**Supplementary Figure 1:** Scheme showing the sequence and position of primers designed for the detection of the different transcript isoforms of *DCP2*. Shown along side in the table and the diagram are the sizes of the expected amplicons.

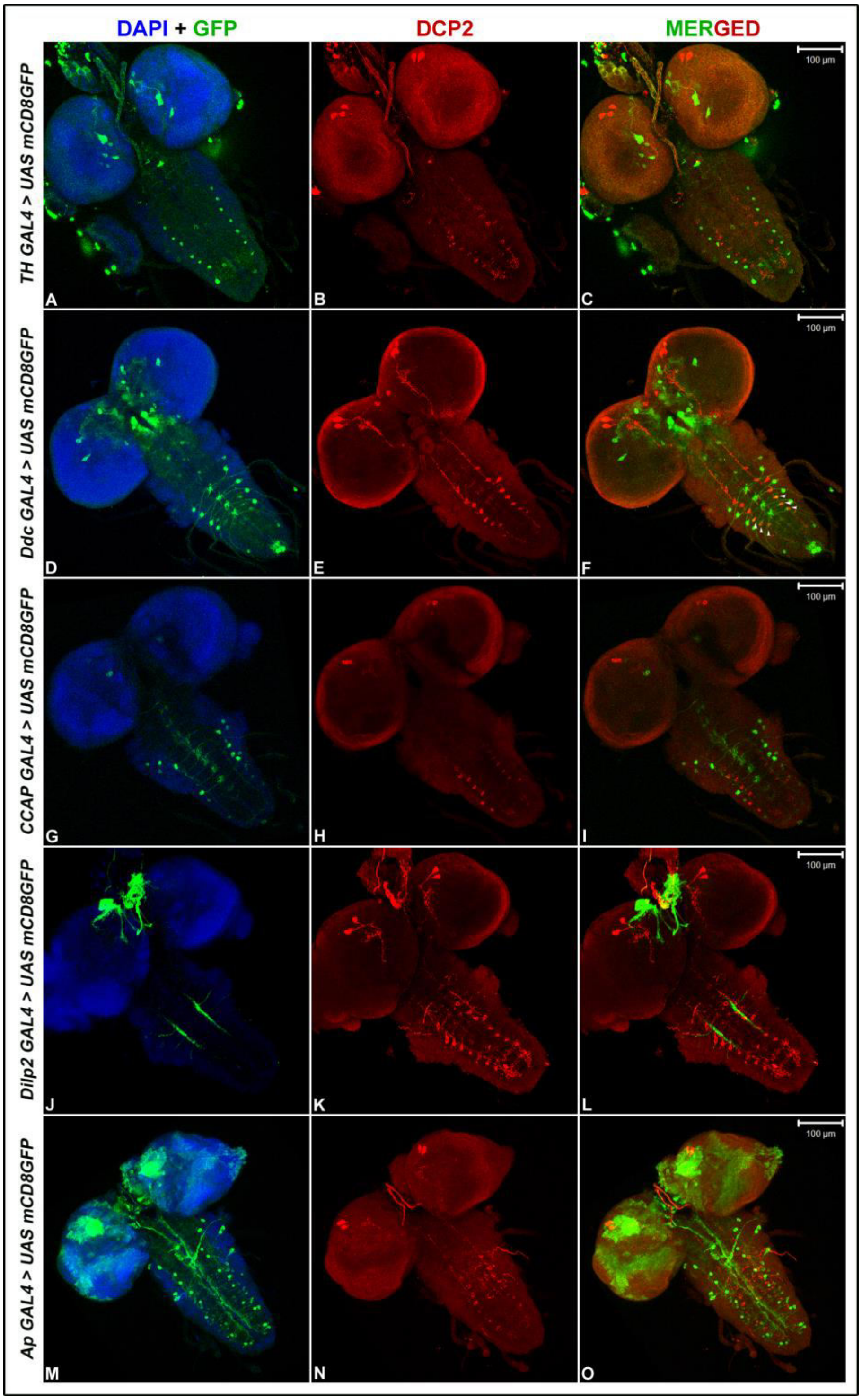


**Supplementary Figure 2:**Confocal projections showing the mapping of DCP2 expressing neurons against the neurons expressing the different biogenic amines Tryptophan hydroxylase (TH; A-C) and Dopamine decarboxylase (Ddc; D-F) , neuropeptides *viz.* Crustacean Cardioactive Peptide (CCAP; G-I), Drosophila Insulin like Peptide-2 (Dilp2; J-L), or the transcription factor Apterous (Ap; M-O).

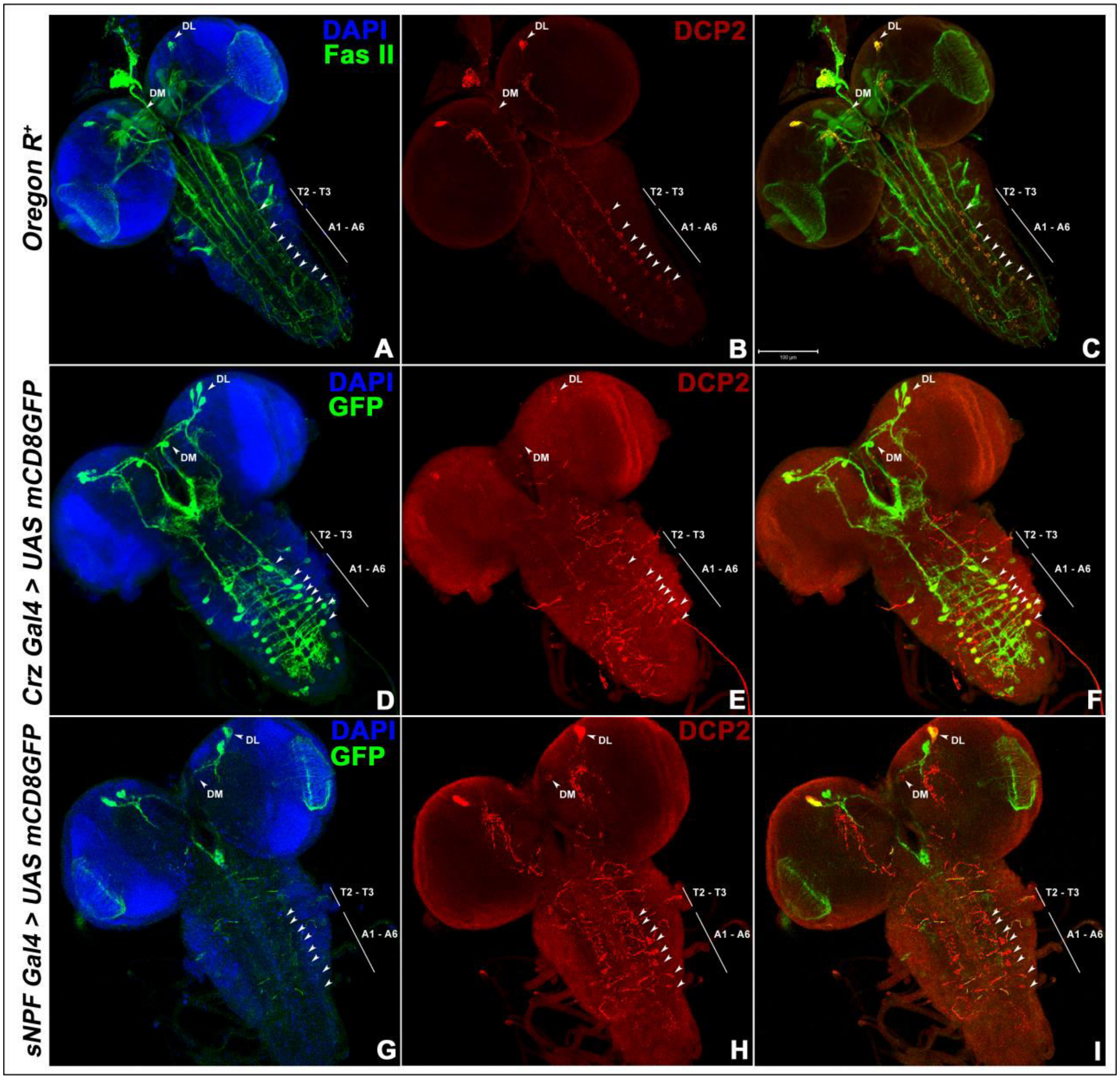


**Supplementary Figure 3:** Confocal Projections showing the overlap of expression of DCP2 with Fasciclin II (A-C) Corazonin (D-F) and sNPF (G-I). DL – dorso-lateral, DM – dorso-medial, T – thoracic, A – abdominal.

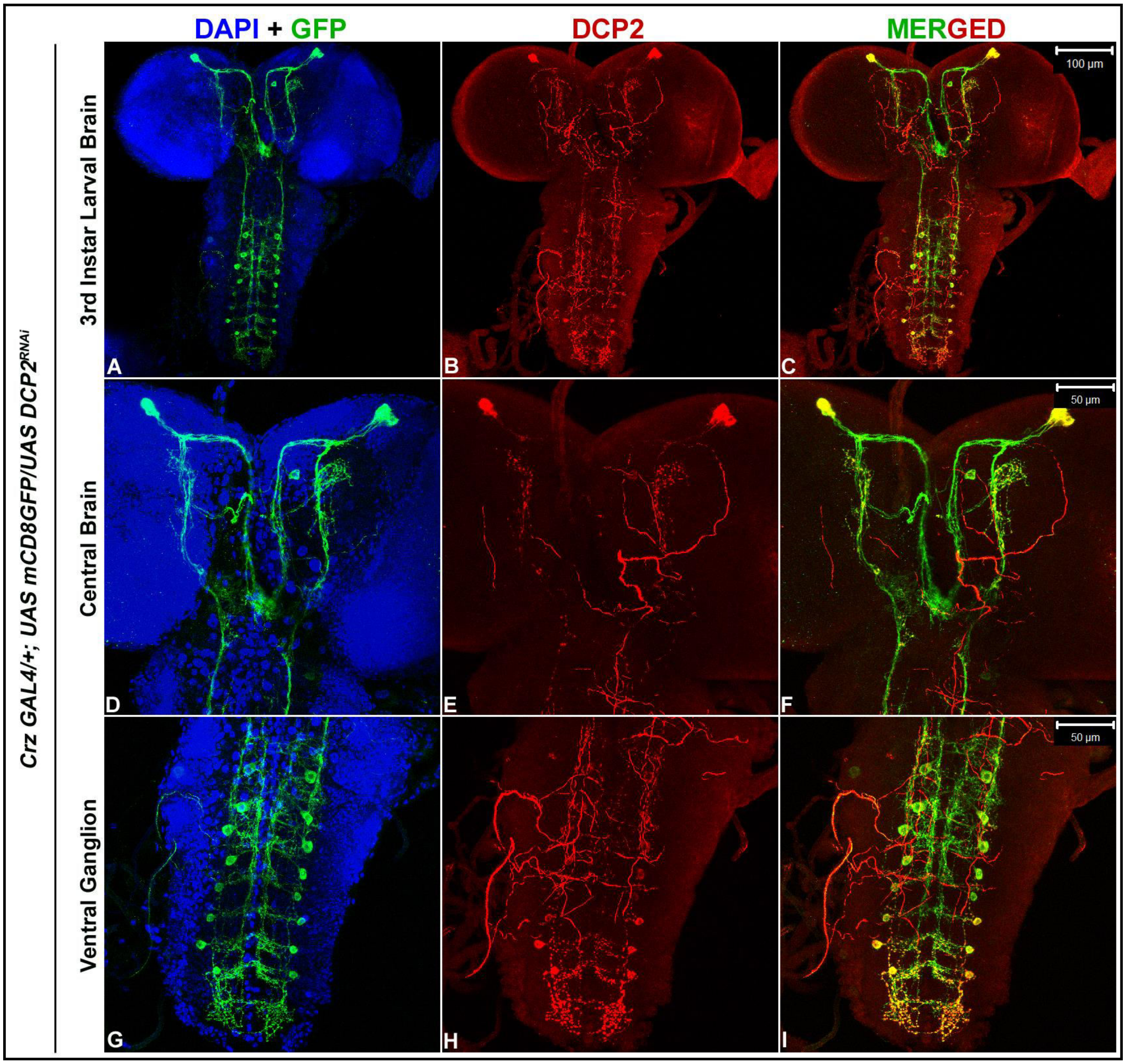


**Supplementary Figure 4:** Confocal projection of brain of 3rd instar larvae showing the expression of DCP2 and GFP in *Crz Gal4>UAS DCP2$^{RNAi}$* larvae. The knock down of *DCP2* does not affect the structure and/or fasciculation of the Crz neurons.
